# RNAseq studies reveal distinct transcriptional response to vitamin A deficiency in small intestine versus colon, uncovering novel vitamin A-regulated genes

**DOI:** 10.1101/798504

**Authors:** Zhi Chai, Yafei Lyu, Qiuyan Chen, Cheng-Hsin Wei, Lindsay M. Snyder, Veronika Weaver, Aswathy Sebastian, István Albert, Qunhua Li, Margherita T. Cantorna, A. Catharine Ross

## Abstract

Vitamin A (VA) deficiency remains prevalent in resource limited countries, affecting over 250 million preschool aged children. Vitamin A deficiency is associated with reduced intestinal barrier function and increased risk of mortality due to mucosal infection. Using *Citrobacter rodentium* (*C. rodentium*) infection in mice as a model for diarrheal diseases in humans, previous reports showed reduced pathogen clearance and survival in vitamin A deficient (VAD) mice compared to their vitamin A sufficient (VAS) counterparts.

**Objectives:** To characterize and compare the impact of preexisting VA deficiency on gene expression patterns in the small intestine (SI) and the colon, and to discover novel target genes in VA-related biological pathways.

**Methods:** VAD mice were generated by feeding VAD diet to pregnant C57/BL6 dams and their post-weaning offspring. RNAseq were performed using the total mRNAs extracted from SI and colon. Differentially Expressed Gene (DEG), Gene Ontology (GO) enrichment, and Weighted Gene Co-expression Network Analysis (WGCNA) were performed to characterize expression and co-expression patterns.

**Results:** DEGs compared between VAS and VAD groups detected 49 SI and 94 colon genes. By GO information, SI DEGs were significantly enriched in categories relevant to retinoid metabolic process, molecule binding, and immune function. Immunity related pathways, including “humoral immune response” and “complement activation” were positively associated with VA in SI. Three co-expression modules showed significant correlation with VA status in SI; these modules contained four known retinoic acid targets. In addition, other SI genes of interest (e.g. *Mbl2*, *Cxcl14*, and *Nr0b2*) in these modules were suggested as new candidate genes regulated by VA. Furthermore, our analysis showed that markers of two cell types in SI, mast cells and Tuft cells, were significantly altered by VA status. In colon, “cell division” was the only enriched category and was negatively associated with VA. Thus, comparison of co-expression modules between SI and colon indicated distinct networks under the regulation of dietary VA and suggest that preexisting VAD could have a significant impact on the host response to a variety of disease conditions.

## Introduction

Vitamin A (VA), retinol and its metabolites, is an essential micronutrient for vertebrates of all ages, from early development throughout the life span. In humans, a nutritional deficiency of VA is both a major cause of xerophthalmia and a risk factor for infectious diseases ^1^. VA deficiency (VAD) is estimated to affect nearly 20% of preschool aged children worldwide ^2^. The impact of VA on child health is well demonstrated by World Health Organization (WHO) data showing that VA supplementation in preschool-age children has reduced diarrhea-related mortality by 33% and measles-related mortality by 50% ^3^.

Several lines of evidence indicate that VA is a pleiotropic regulator of gut immunity ^4^. Retinoic acid (RA), the most bioactive metabolite of VA, has been shown to influence the differentiation ^5^ and gut homing of T cells ^6^, secretory IgA production ^7^, and the balance of different innate lymphoid cell (ILC) populations ^4^. Regarding intestinal epithelial cells, VA affects both cell differentiation and function of the intestinal epithelium ^8^. In murine models, VA deficiency has been associated with abnormality in goblet cell and Paneth cell ratios and activities ^9^, as well as weakened intestinal barrier integrity ^10^. In sum, VA is a versatile regulator of both innate and adaptive immunity in the gut.

Preexisting VA status has been shown to affect the immune response in several animal models of gut infection, including infection with *Citrobacter rodentium* (*C. rodentium*), a gram negative natural mouse intestinal pathogen that mimics enteropathogenic *Escherichia coli* infection in humans ^11^. Host VA status significantly affects the host response to *C. rodentium*, for vitamin A sufficient (VAS) mice cleared the infection and survived, whereas vitamin A deficient (VAD) mice suffered a 40% mortality rate and those mice that survived the infection became non-symptomatic carriers of *C. rodentium* ^12^.

The initial VA status of mice at the time of infection could be an important factor in their ability to respond to gut infection. Previous studies did not comprehensively examine gene expression in the small intestine (SI) or colon of mice according to their VA status. In the present study, we sought to determine key genes that may mediate the divergent host responses between VAS and VAD mice, by comparing intestinal gene expression in naïve VAS and VAD mice prior to *C. rodentium* infection. Although *C. rodentium* infects primarily the colon ^13^, the SI is also relevant because it is a major site of retinol uptake into enterocytes, formation of retinyl esters for absorption into the body, and local conversion to metabolites such as RA ^14^. In the steady state, retinol concentrations are significantly higher in the SI and mesenteric lymph nodes compared with most other tissues, excepting the liver, the major storage site of VA ^15^. Intestinal epithelial cells and dendritic cells (DCs) in SI are producers of RA and the intestinal RA concentration generally follows a gradient from proximal to distal SI, correlating with the imprinting ability of resident DCs ^15^. Therefore, both SI and colon were of interest for our present study in which we have used differential expression and co-expression analyses to characterize the effect of VA on the transcriptomic profiles in these two organs. Additional experimental groups and analyses focused on the effect of infection and interaction with VA status were also performed, and will be reported separately. Here, we have focus on the effect of VA status alone as a presumptive predetermining risk factor for the host response to gut infection.

## Results

### Model validation: Serum retinol is lower in VAD mice than VAS mice

VAS and VAD mice were generated by feeding diets that were either adequate or deficient in VA to pregnant mice and to their offspring for eight to ten weeks. The VA status of our animal model was validated by determining serum retinol concentrations at the end of the SI study ^16^. Serum retinol levels were significantly lower in VAD vs. VAS mice (*P*<0.001) (Fig. 1), showing that VA deficiency was successfully induced in our model and enabling us to further analyze the effect of VA status on intestinal gene expression.

**Fig. 1.**
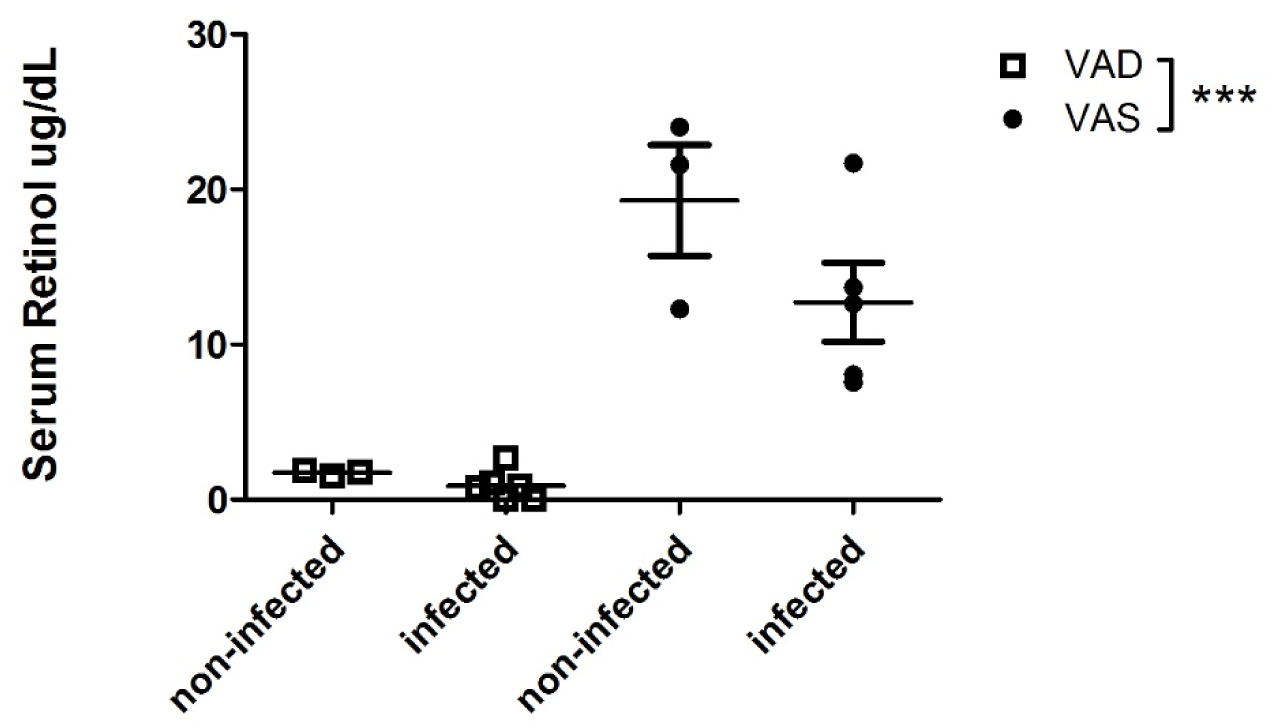
VAD mice have lower serum retinol levels than VAS mice. Serum retinol concentrations were measured by UPLC on post-infection day 5. Values are the mean ± SEM of n=3-6 mice per group. Two-way ANOVA test. *** *P*<0.001. **Abbreviation:** vitamin A (VA), vitamin A deficient (VAD), vitamin A sufficient (VAS), Ultra Performance Liquid Chromatography (UPLC), standard error of the mean (SEM), analysis of variance (ANOVA).

### Distribution of differentially expressed genes in SI and colon and their co-expression

We elected to use whole tissue RNAseq analysis to provide a comprehensive exploratory view of the expression profiles in both SI and colon ^17^. Initial mapping for the SI revealed 24,421 genes, which were subsequently screened for low expression transcripts and normalized by DESeq2 (Fig. 2). Among the 14,368 SI-expressed genes, 49 differentially expressed genes (DEGs) differed with VA status, and 9 co-expression modules were identified. Similarly, among the 15,340 genes mapped in the colon dataset, 94 DEGs differed with VA status and 13 co-expression modules were identified. Of these, 14,059 genes were shared between the two datasets and these were used for the module preservation analysis (Fig. 2).

**Fig. 2.**
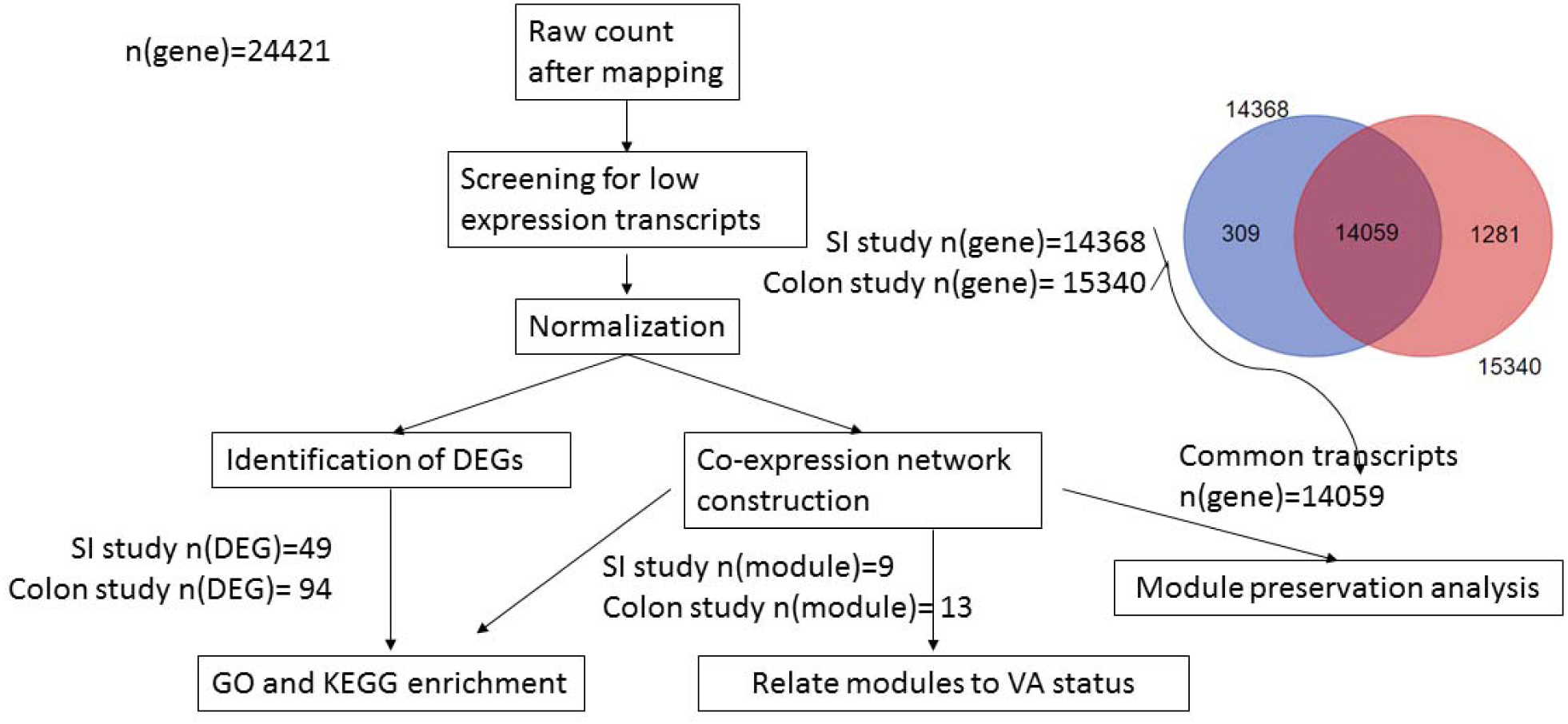
Overview of the results of bioinformatics analyses. Initial mapping identified 24,421 genes. After data pre-processing, 14,368 genes in the SI study and 15,340 genes in the colon study were maintained in the two datasets. Starting from the 14,368 SI-expressed genes, 49 DEGs corresponded to VA effect, and 9 co-expression modules were identified. Similarly for the colon study, 94 DEGs and 13 modules were identified among the 15,340 colon-expressed genes in the colon study. A total of 14,059 genes were shared between the two datasets and were used to for the module preservation analysis.

### Identification of DEGs

Of the 49 DEGs in the SI that differed significantly by VA status, 18 were upregulated (higher in VAS than VAD, Supplementary Table 1) and 31 were downregulated (lower in VAS than VAD, Supplementary Table 2). A heatmap of the DEGs in the SI study is shown in Supplementary Fig. 1. A volcano plot of the SI DEGs (Fig. 3a) shows that genes with the largest negative or positive fold changes as well as highest statistical significance included *Cd207*, *Mcpt4*, and *Fcgr2b*, which were lower in VAS than VAD mice, and *Isx*, *Cyp26b1*, *Mmp9*, and *Mbl2*, which were higher in VAS than VAD mice.

**Fig. 3.**
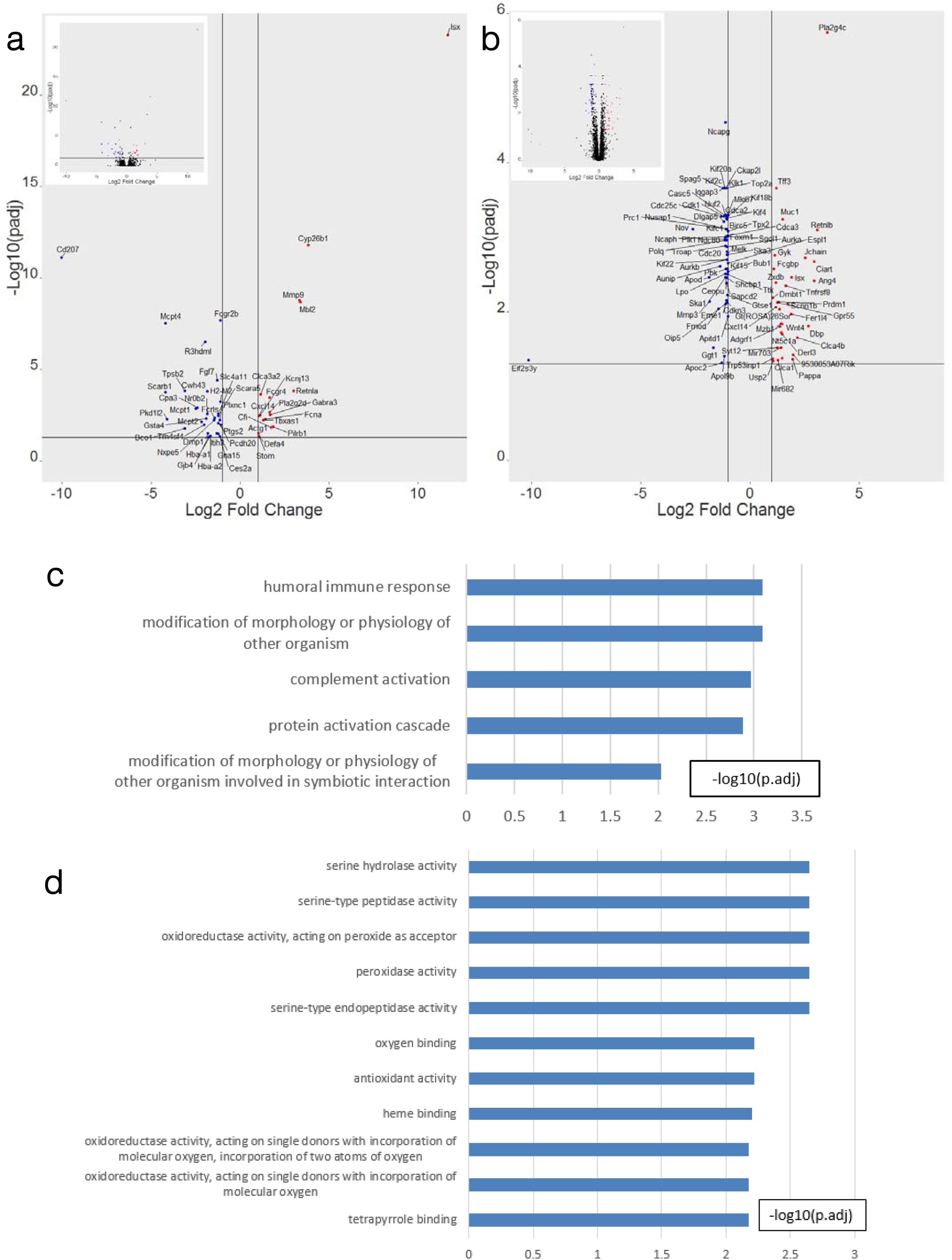
Transcriptional changes in the intestines corresponding to VA status. **a**, **b**, Volcano plots of DEGs in SI (**a**) and in colon (**b**). Red: DEGs upregulated in VAS group comparing with VAD group; Blue: DEGs downregulated in VAS group comparing with VAD group; Black: non-DEGs. Criteria: |Fold Change|=2 is marked out by two black vertical lines x=1 and x=-1. adjp<0.05 is marked out by a horizontal black line y= - Log10(0.05). **c**, **d**, GO enrichment terms upregulated (**c**) or downregulated (**d**) by VA in SI. Criteria: p.adj<0.05. **Abbreviations:** padj: adjusted *P* value according to DESeq2, differentially expressed gene (DEG), SI (small intestine), VAS (vitamin A sufficient), VAD (vitamin A deficient), Gene ontology (GO), p.adj (BH-adjusted *P* value from the hypergeometric test conducted by ClusterProfiler).

Of the 94 DEGs in the colon, 34 were more highly expressed in VAS than VAD mice (Supplementary Table 3) and 60 genes were of lower expression (Supplementary Table 4). The heatmap for colon (Supplementary Fig. 2) and volcano plot (Fig. 3b) show that some of the most significantly upregulated DEGs are also well-known players in goblet cell activities, for example, *Tff3* and *Muc1* ^18^, known as two goblet cell markers; and *Clca1* and *Clca4b*, known to regulate mucus production and neutrophil recruitment ^19^. Several genes with known regulatory functions were among the most strongly regulated in our datasets: for example, three transcription factors (TFs), *Isx*, *Prdm1*, and *Zxdb* were more highly expressed in VAS mice, while *Klk1*, a mucus regulator and *Foxm1*, a TF, were downregulated.

### Gene ontology (GO) and KEGG enrichment of DEGs

We next performed enrichment analysis separately on all DEGs, upregulated DEGs, and downregulated DEGs to compare the VAS vs VAD condition in the two datasets (Fig. 2). Not surprisingly, the 49 SI DEGs were significantly enriched in “retinoid metabolic process” (Supplementary Fig. 3), including in VAD mice a higher expression of *Bco1*, a beta-carotene cleavage enzyme, and in VAS mice a higher expression of *Cyp26b1*, a retinoic acid degrading enzyme, and *Isx*, a master regulator and repressor of *Bco1* and other retinoid-pathway genes. The 18 upregulated SI DEGs were significantly enriched in five Biological Processes (Fig. 3c), whereas the 31 downregulated SI DEGs were significantly enriched in 10 categories of Molecular Function (Fig. 3d).

In the colon, downregulated DEGs (*n*=60) in VAS vs VAD mice were significantly enriched in “Cell Division” (GO term) and “Cell Cycle” (KEGG term, mmu04110), while upregulated colon DEGs (*n*=34) were not enriched in either GO or KEGG pathways.

### WGCNA modules significantly correlated with VA status

We next applied Weighted Gene Co-expression Network Analysis (WGCNA) as a systems biology approach to understanding gene networks in contrast to individual genes (Fig. 2). In the SI, setting a soft threshold power β to 7 (Fig. 4a), WGCNA identified nine modules with module size (MS) ranging from 202 to 5,187 genes (Supplementary Table 5). These modules in the SI network will be referred to henceforth by their color labels [e.g., SI(Color)], which is a standard approach wherein the colors are merely labels arbitrarily assigned to each module. Modules labeled by colors are depicted in the hierarchical clustering dendrogram in Fig. 4b. The Grey module was used to hold all genes that do not clearly belong to any other modules. The gene expression patterns of each module across all samples are demonstrated as heatmaps (Supplementary Fig. 4).

**Fig. 4.**
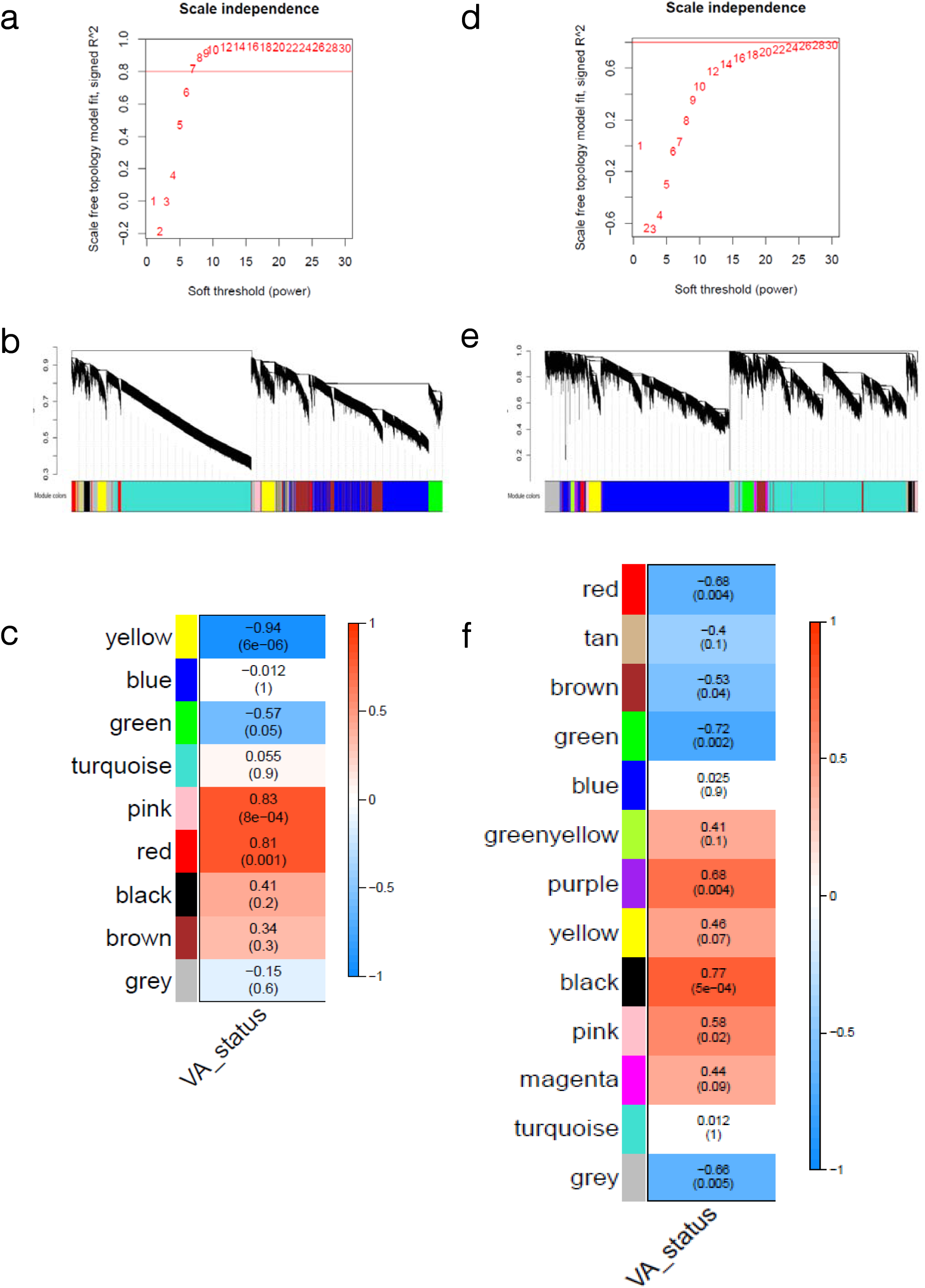
Co-expression network construction in the intestines via WGCNA. **a**, **b**, **c**, Results from SI study. **d**, **e**, **f**, Results from colon study. **a**, **d**, Soft threshold power beta were chosen for SI (**a**) and colon (**d**) networks according to the scale-free topology criteria (horizontal red lines). **b**, **e**, Clustering dendrograms of co-expression modules in SI (**b**) and colon (**e**). Transcripts are represented as vertical lines and are arranged in branches according to topological overlap similarity using the hierarchical clustering. The dynamic treecut algorithm was used to automatically detect the nine highly connected modules, referred to by a color designation. Module color designation are shown below the dendrogram. **c**, **f**, Correlation of module eigengenes with VA status in SI (**c**) and colon (**f**) networks. Upper numbers in each cell: Correlation coefficient between the module and the trait. Lower numbers in parentheses: *P* value of the correlation. Significant correlations (*P* <0.01) were obtained between VA status and the SI(Yellow), SI(Pink), SI(Red) (**c**), Colon(Black), Colon(Green), Colon(Purple), and Colon(Red) (**f**) modules. **Abbreviation:** weighted gene co-expression network analysis (WGCNA), small intestine (SI), vitamin A (VA).

To determine if any of the nine modules of co-expressed SI genes were associated with VA status, we tested the correlations between the module eigengenes and VA status (Fig. 4c). Eigengene is defined as the first principal component of a given module and may be considered as a representative of the gene expression profiles in that module ^20^. Three modules were significantly correlated with VA status: the SI(Yellow), SI(Pink), and SI(Red) modules, with *P* values of 6 x 10^-6^, 8 x 10^-4^ and 0.001, respectively.

Among these three modules, Yellow module is the largest (MS=924), with a correlation coefficient of −0.94 with VA status (Fig. 4c), thus, this module contains transcripts that were lower in the SI of VAS mice vs. their VAD counterparts (Supplementary Fig. 4). According to ClusterProfiler, the SI(Yellow) module is functionally enriched in genes for adaptive immunity and cytokine activity (Supplementary Table 5). The SI(Yellow) module contains known key players in RA metabolism (*Bco1*, *Scarb1*, *Gsta4*, *Aldh1a2*, and *Aldh1a3*), as well as potential RA targets (*Rdh1*, *Aldh3a2*, *Aldh7a1*, and *Aldh18a1*), along with several immune cell markers (e.g. *Cd1d1*, *Cd28*, *Cd44*, *Cd86*, and *Cd207*).

The SI(Pink) module was positively correlated with VA status, however no functional annotation was enriched in this module.

In contrast, the SI(Red) module was positively correlated with VA status and enriched for functional categories involved in innate immune response (Supplementary Table 5). SI(Red) module includes genes previously reported as VA targets *Isx*, *Ccr9*, and *Rarb*, as well as some immunological genes of interest, e.g., *Lypd8*, *Cxcl14*, *Mmp9*, *Mbl2*, *Noxa1*, *Duoxa2*, *Muc13*, *Itgae*, *Itgam*, *Madcam1*, *Il22ra2*, *Pigr*, *Clca1*, *Clca3a2*, and *Clca4b*, as the product of these genes could mediate the effects of VA status on infection. Overall, comparing VAS and VAD groups, our analysis suggests that innate immune activity is stronger, while adaptive immune response is weaker in the VAS SI.

In order to visualize the co-expression relationships within the modules of interest, for each of these three modules the top 50 genes selected by connectivity are depicted as networks in Fig. 5. TFs are usually more central in the network, examples include *Nr0b2* (Fig. 5a), *Isx* (Fig. 5b), and *Smad4* (Fig. 5c), where *Nr0b2* and *Isx* were also identified as DEGs.

**Fig. 5.**
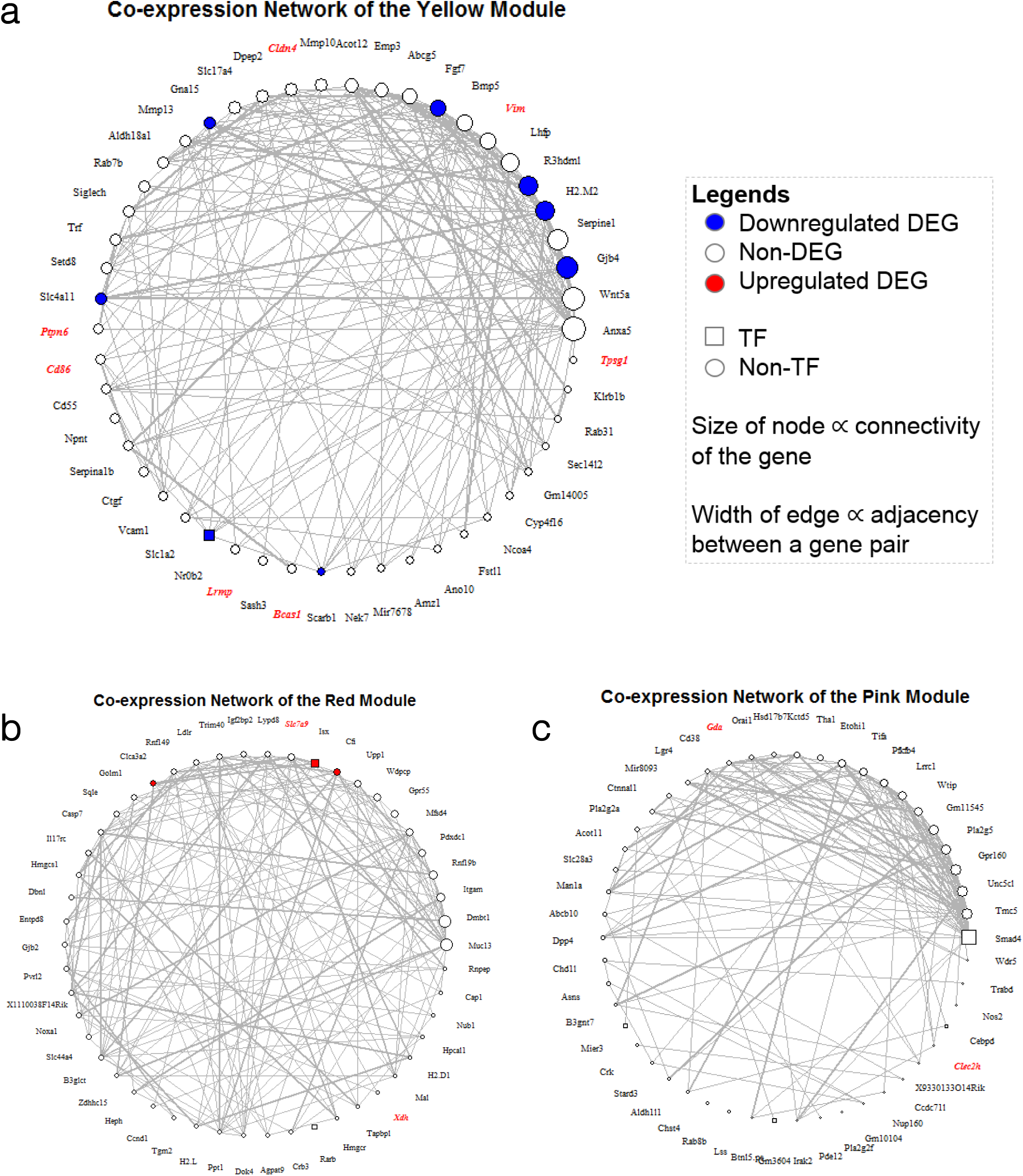
Visualization of the co-expression network in three modules of interest in SI. **a**, SI(Yellow), **b**, SI(Red), and **c**, SI(Pink). In each of the modules of interest, the top 50 genes ranked by connectivity are depicted using igraph in R. Each node represents a gene. The size of the node is proportional to the connectivity of the gene. The shape of the nodes is coded squares = TFs, circles = non-TFs. Color of the nodes denote: Blue = upregulated DEGs in VAS condition, Red =downregulated DEGs. Font: red bold italic = cell type markers. Thickness of lines between nodes: proportional to the adjacency value of the gene pair. In order to improve network visibility, adjacencies equal to 1 or lower than 0.7 were omitted. **Abbreviations**: SI (small intestine), TF (transcription factor), VAS (vitamin A sufficient).

Using the same WGCNA analysis on colon transcriptomes (with soft threshold power β of 26, Fig. 4d), 13 modules were identified and color-coded [referred by Colon(Color), Fig. 4e and Supplementary Fig. 5], with MS ranging from 87 to 5,799 genes (Supplementary Table 6). Four modules, Colon(Black), Colon(Green), Colon(Purple), and Colon(Red) differed significantly with VA status, with *P* values of 5 x 10^-4^, 0.002, 0.004, and 0.004, respectively (Fig. 4f). Among these four modules, Colon(Green) was enriched for “cytoplasmic translation,” “ribosome biogenesis,” and “ncRNA processing,” while Colon(Red) was enriched for “anion transmembrane transporter activity” (Supplementary Table 6).

### Associations with Cell Markers

To further query whether the expression pattern of each module was mainly driven by a regulatory effect of VA on cell differentiation, a cell type marker enrichment was performed. This test identified four SI modules that significantly overlap with the list of SI cell markers derived from the CellMarker database ^21^ (Fig. 6). The SI(Yellow) module, which as noted above is negatively correlated with VA (Fig. 4c), contains 24 Tuft cell markers, reaching a Bonferroni corrected *P* value (CP) of 4 x 10^-7^. This suggested a negative regulation of VA on the Tuft cell population. The SI(Green) module also significantly overlapped with Tuft cell markers (CP=0.01). The SI(Turquoise) module, the largest module with MS=5187 genes (Supplementary Table 5), was significantly enriched for markers of goblet cells (CP=1 x 10^-6^) and Paneth cells (CP=0.001).

**Fig. 6.**
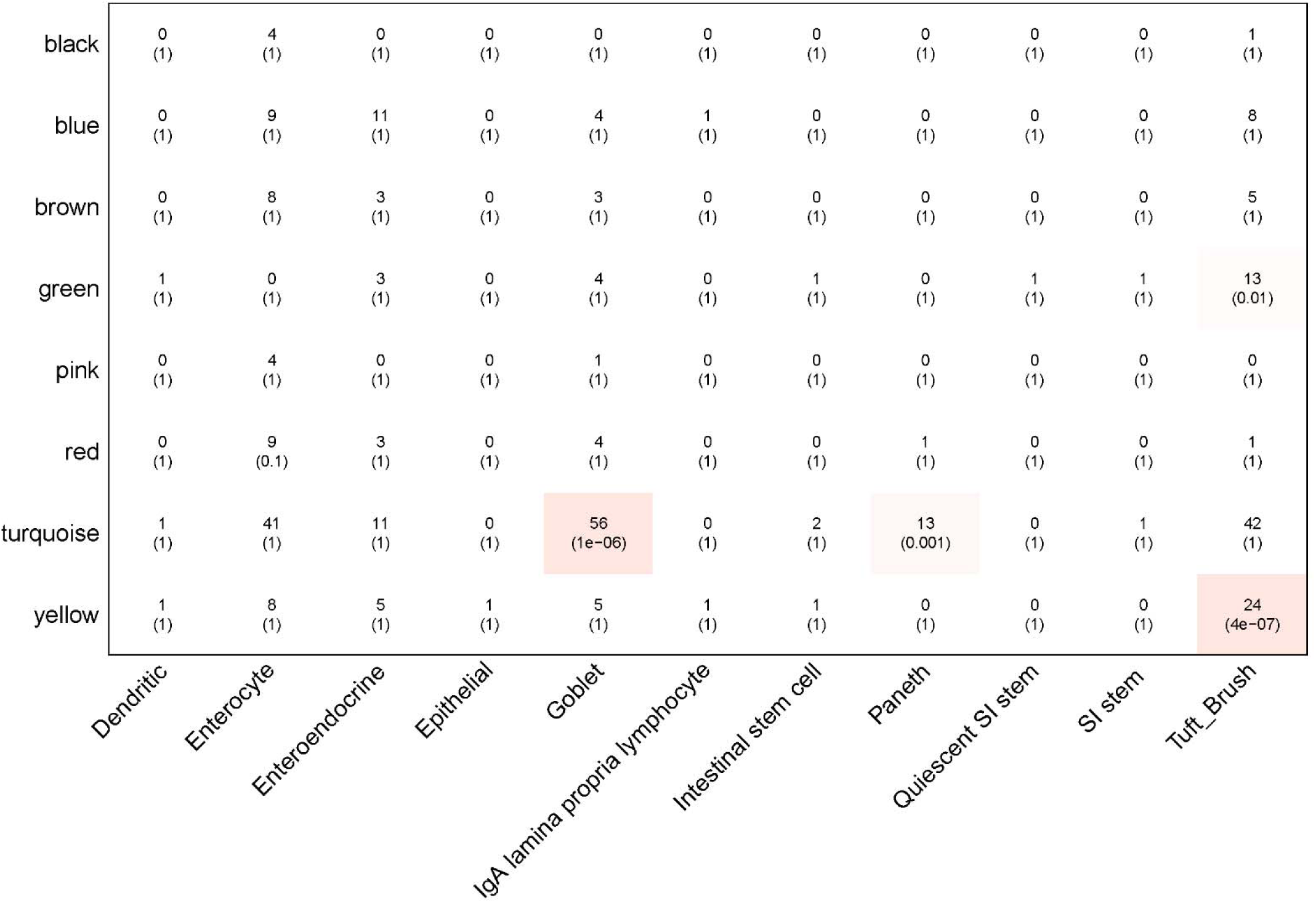
Overlap between module genes and SI cell markers. The cell type marker lists of SI were obtained from CellMarker, a manually curated database. Upper numbers in each cell: Number of probes shared between the designated module (y axis) and the cell marker list (x axis). Lower numbers in parentheses: Bonferroni corrected *P* value (CP) of the one-sided hypergeometric test, which evaluates if the gene list of a module significantly over-represents any cell type markers. Criteria: CP<0.05. Color shade of each cell is according to –log10(CP). Four significant overlaps were highlighted with red shades.

### Module preservation analysis suggests that gene networks controlled by VA are poorly preserved between SI and colon

To determine whether the modules identified as being significantly correlated with VA in SI have a similar module structure in the colon, we performed module preservation analysis using 14059 genes that were robustly expressed in both datasets (Fig. 2). Network constructions were plotted for SI and colon separately. The modules from this analysis are referred as mpSI(Color) and mpColon(Color). Module preservation analysis was performed on mpSI and mpColon networks. For all three mpSI modules that were significantly correlated with VA status (Red, Pink, and Yellow), we found only “weak to moderate” evidence of preservation in the mpColon network according to Zsummary (Supplementary Table 7). In addition, mpSI Red and Pink, two modules associated with VAS status (Fig. 4c), overlapped merely with the mpColon modules broken up into Black1 and Black2 (Supplementary Fig. 6) instead of an intact mpColon(Black) module, which was originally also associated with VAS status (Fig. 4f). mpSI(Yellow), another SI module significantly correlated with VA status (Fig. 4c), had a lower preservation score than the mpSI (Red) and mpSI(Pink) modules (Supplementary Table 7), and did not significantly overlap with any VA-related modules (Black, Green, Purple, or Red) in the mpColon network (Supplementary Fig. 6).

## Discussion

### Differential expression analysis confirmed known targets under the control of VA in SI and validated the nutritional model

The approach we have taken in this study was to use rapidly frozen intact tissue to capture a “bird’s eye” comprehensive view of an entire organ, SI and colon individually, in physiologically normal mice subjected to only dietary treatment as a means to represent VA adequacy and deficiency in human populations. The use of intact tissue eliminated the selective loss or enrichment of cells and genes within, such as could occur during intestinal cell isolation protocols ^22^. On the other hand, a limitation of our approach is that signals from genes significantly regulated by VA but residing within low-abundance cell types might have been overwhelmed and thereby missed. We first found that of the 49 DEGs differing by VA status in SI there was a strong enrichment in “retinoid metabolic process” driven by three DEGs: *Cyp26b1* (Fold Change=14.1), *Isx* (Fold Change=3,231), and *Bco1* (Fold Change= −8.7) in VAS vs. VAD mice (Supplementary Table 1 and 2). RA is known to induce the transcription of CYP26B1, an enzyme that determines the magnitude of cellular exposure to RA by inactivating intracellular RA, via nuclear receptor protein binding to multiple retinoic acid response elements in the gene’s promoter region ^23^; *Cyp26b1* has been shown to be dynamically regulated in rodent liver ^24^, lung ^25^, and SI ^26^. RA also induces the expression of ISX ^27^, a transcriptional repressor for both BCO1, which catalyzes ß-carotene cleavage, and SCARB1, a receptor for carotene uptake, in the intestine ^28^. Thus, the sum of the upregulation of *Cyp26b1* and *Isx*, as well as the downregulation of *Scarb1* and *Bco1* in the VAS SI, confirms the well-established negative feedback regulation of dietary VA on the intestinal RA concentration.

At the same time, a series of immunological genes previously reported to be regulated by VA, albeit in a variety of tissues, were also identified in the SI dataset, including *Mmp9*, *Cd207*, and enzymes produced by mast cells. The SI contains the body’s largest population of immune cells ^29^, with many cell types present, and thus finding genes in this category in our study was not unexpected. However, finding genes in this category that are sensitive to VA status may be informative for designing future studies. MMP9 is an extracellular matrix-degrading enzyme ^30^ that plays important immunological roles in macrophages and DCs ^31^, wherein *Mmp9* expression is induced by RA. In mouse myeloid DCs, chromatin immunoprecipitation assays showed that *Mmp9* expression was enhanced by RA through a transcriptional mechanism involving greater RAR-alpha promoter binding and chromatin remodeling ^32^. CD207, also known as langerin, is a marker of the langerin^+^ DCs, a cell type normally only abundant in skin but not in the gut ^33^. However, it has been reported that RA deficiency can change DC subtypes in the intestine and dramatically elevate langerin^+^ DC numbers ^34^, which is in concordance with our SI study, where *Cd207* expression was significantly higher in the VAD group (Fig. 3a and Supplementary Table 2). Mast cell proteases (MCPT) 1, 2, 4, and 6 (also known as TPSB2 or tryptase beta 2) were uniformly downregulated in the VAS group, an observation in line with previous reports that retinol and RA inhibit the growth and differentiation of human leukemic mast cells (HMC-1) ^35^, and the mast cells derived from peripheral blood or cord blood samples ^36^. In addition, peritoneal mast cell numbers were also reported to be 1.5 times higher in VAD rats ^37^. Overall, both predictable observations based on previous analysis of the SI, and observations in the SI that are congruent with the effects of VA and RA shown in other cell types, lend support to the strong impact that VA nutrition has on retinoid homeostasis and immune-related genes in the SI.

### Potential new target genes regulated by VA in SI

Our analysis also uncovered several genes that were significantly regulated by VA status that may be targets for further study in the context of gut infections. One of these is *Mbl2*, an active player in gut immunity, the expression of which may be controlled by VA in SI (Fig. 3a). Mannose-binding lectin (MBL) is an important pattern-recognition receptor in the innate immune system ^38^, and MBLs are protective against gut infection by selectively recognizing and binding sugars presented on the pathogens ^39^. This binding can neutralize the pathogen and inhibit infection by complement activation through opsonization or through the lectin pathway. Low levels of MBL have been related to increased susceptibility to several pathogens such as *Neisseria meningitidis, Staphylococcus aureus*, and *Streptococcus pneumoniae* ^40^. In regards of the transcriptional regulation. SI is a predominant site of extrahepatic expression of MBL. Allelic variants of *Mbl2* have been associated with deficiencies in MBL protein production, which may increase the risk of mother-to-child HIV transmission ^41^. However, interestingly, among infants with *Mbl2* variants, VA plus β-carotene supplementation partially counteracted the increased risk of transmission ^42^. Currently, no mechanistic study has investigated how VA affects *Mbl2* gene expression; however, since *Mbl2* is an upregulated DEG ranked high in terms of both statistical significance and Fold Change (Fig. 3a and Supplementary Table 1), we postulate that VA may be an inducer of *Mbl2* gene expression. If so, studies of this gene may provide another line of evidence that VA participates in the regulation of the innate mucosal immunity.

Another candidate identified in our analysis is the *Nr0b2* (nuclear receptor subfamily 0, group B, member 2), encoding the TF small heterodimer partner (SHP). SHP is an atypical nuclear receptor that binds to a variety of nuclear receptors ^43^. In liver, SHP plays an important role in controlling cholesterol, bile acid, and glucose metabolism. Hepatic *Nr0b2* is retinoid-responsive both *in vitro* and *in vivo* ^44, 45^. In the hepatocyte cell line AML12, *Nr0b2* was upregulated dose-dependently by RA ^44, 46^. However, in our SI study, *Nr0b2* was downregulated by VA (Fig. 3a and Supplementary Table 2). This may be due to organ specificity in the transcriptional regulation on *Nr0b2* gene, and/or may reflect the different roles played by liver and SI in cholesterol metabolism. *Nr0b2* ranks relatively high regarding connectivity in our SI(Yellow) module (Fig. 5a), which is in line with its annotation as a TF.

### VA as a potential regulator of the SI Tuft cell population

An interesting observation that emerged from using the CellMarker database to identify cell type-selective genes in the SI dataset was the significance enrichment of Tuft cell markers in the SI(Yellow) module (Fig. 6). As this module is negatively correlated with VAS status, this enrichment suggests that the SI Tuft cell population may be more abundant under VAD status (Fig.4c and Supplementary Fig. 4). Tuft cells normally represent only about 0.5% of the epithelial cells in the murine small and large intestines, with a slightly more prevalent distribution in the distal SI ^47^. As Tuft cells are a relatively newly identified secretory cell type ^48^, they have not yet been well studied. We found no literature on this cell type with respect to VA status. A speculation is that VAD status is associated with higher numbers of ILC2 cells, which stimulate Tuft cell differentiation. VA is known to regulate the balance between ILC2s and ILC3s in the intestinal mucosa ^4^. VA deficiency has been shown to suppress ILC3 cell proliferation and function, and reciprocally to promote ILC2s ^4^. IL-13 is one of the ILC2 cytokines that can drive the differentiation of epithelial precursors towards the Tuft lineage ^47^. One of the advantages of the co-expression network analysis used in our study is that cell type-specific information may be derived through the analysis of whole intact tissues, without the isolation of homogeneous populations of cells ^22, 49^. Given the small number of Tuft cells, isolation may prove difficult but approaches such as single-cell RNAseq, laser capture microdissection, and immunohistochemistry may be useful in the future. Although markers of goblet and Paneth cell were significantly enriched in SI(Turquoise) module, this module did not show significant correlation with VA status (Fig. 4c).

### VA differentially regulates SI and colon transcriptomes

In general, DEGs in the SI were fewer in number but more highly regulated by VA than those in the colon. The 49 DEGs in SI not only included cell markers (Supplementary Table 2), but also genes in immunological and retinoid metabolic pathways (Fig. 3 c, d and Supplementary Fig. 3). Compared with the SI, the colon was enriched for fewer categories that differed by VA status. Of 94 colon DEGs, their GO terms related to cell division, which suggests that the primary transcriptional role of VA in colon is related to inhibition of cell proliferation and induction of epithelial differentiation, two widely applicable functions of RA. Together with the upregulation of goblet cell markers (*Tff3*, *Muc1*, *Clca1*, and *Clca4b*), our results are consistent with a differentiation-promoting effect of VA in the colon (Supplementary Table 3). Additionally, even though *Isx* was a DEG responsive to VA in both SI and colon, we did not find *Bco1* and *Scarb1* to be differentially expressed in the colon, in line with the processes of carotenoid digestion and absorption taking place primarily in SI. Other than *Isx* and *Cxcl14*, a potent chemoattractant for immune cells ^50^, no other genes in our studies were induced by VA in both organs.

Additionally, co-expression network analysis of the VA-correlated modules revealed very distinctive module structures between the SI and colon. Although most genes were commonly expressed in both organs (*n*=14,059, Fig. 2), the preservation and overlap of VA-correlated modules between the two organs were likely to be insignificant as judged from the Zsummary score (Supplementary Table 7) and the lack of counterpart modules (Supplementary Fig. 6). In sum, the networks under the influence of VA in SI appeared poorly preserved in colon. This is a further proof that although SI and colon are consecutive organs with similar developmental origins, their biological functions, especially their roles in VA metabolism, are very distinct from each other.

## Conclusions

Taken together, the results of our analyses suggest the following: DEGs corresponding to VA effect are present in both the SI and colon. In SI, DEGs altered by VA are mainly involved in the retinoid metabolic pathway and immunity-related pathways. Our analysis uncovered novel target genes (e.g. *Mbl2*, *Cxcl14*, and *Nr0b2*) and moreover suggested that certain cell types—mast cells based on differential expression analysis and Tuft cells based on marker analysis—are under the regulation of VA, and therefore appear to be promising targets for further study. With regard to the colon, VA has fewer large effects but appears to play an overall suppressive role on cell division while promoting cell differentiation. Finally, the comparison of co-expression modules between SI and colon indicates distinct regulatory networks with few overlaps under the control of VA. Collectively, these data provide a new framework for further investigations of VA-regulated cell differentiation in the SI and colon, and for studies of the impact of infection on genes regulated by VA.

## Methods

### Animals

C57BL/6 mice, originally purchased from Jackson Laboratories (Bar Harbor, ME, USA), were bred and maintained at the Pennsylvania State University (University Park, PA, USA) for experiments. Mice were exposed to a 12 hour light, 12 hour dark cycle with *ad libitum* access to food and water. All animal experiments were performed according to the Pennsylvania State University’s Institutional Animal Care and Use Committee (IACUC) guideline (IACUC # 43445). Vitamin A deficient (VAD) and vitamin A sufficient (VAS) mice were generated as previously described ^12, 16, 51, 52^. Briefly, VAS or VAD diets were fed to pregnant mothers and the weanlings. VAS diet contained 25 μg of retinyl acetate per day whereas VAD diet did not contain any VA. At weaning, mice were maintained on their respective diets until the end of the experiments. The SI study lasted 5 days following *C. rodentium* infection, whereas the colon study ended during the peak of infection (post-infection day 10). Serum retinol concentrations were measured by UPLC (Ultra Performance Liquid Chromatography) to verify the VA status of experimental animals.

### UPLC serum retinol quantification

Serum samples were saponified and quantified for total retinol concentration by UPLC ^53^. Briefly, serum aliquots (30–100 μl) were incubated for 1 h in ethanol. Then 5% potassium hydroxide and 1% pyrogallol were added. After saponification in a 55°C water bath, hexanes (containing 0.1% butylated hydroxytoluene, BHT) and deionized water were added to each sample for phase separation. The total lipid extract was transferred and mixed with a known amount of internal standard, trimethylmethoxyphenyl-retinol (TMMP-retinol). Samples were dried under nitrogen and re-constituted in methanol for UPLC analysis. The serum total retinol concentrations were calculated based on the area under the curve relative to that of the internal standard.

### Statistical analysis

Statistical analyses were performed using GraphPad Prism software (GraphPad, La Jolla, CA, USA). Two-way analysis of variance (ANOVA) with Bonferroni’s *post-hoc* tests were used to compare serum retinol levels. *P*<0.05 was used as the cut off for a significant change.

### Tissue collection and RNA extraction

Tissue were collected at the end of each study. During tissue collection, the dissected colons were first cut open to remove fecal contents. Both lower SI and colon tissues were snap frozen in liquid nitrogen immediately upon tissue harvest. In the SI study, lower SI tissue was used for total RNA extraction. In the colon study, the whole colon tissue free of colonic content was used to extract total RNA. For both tissues, total RNA extractions were performed using the Qiagen RNeasy Midi Kit according to the manufacturer’s protocol. RNA concentrations were measured by Nanodrop spectrophotometer (NanoDrop Technologies Inc., Wilmington, DE, USA). TURBO DNA-free kit (Ambion, Austin, TX, USA) were used to remove genomic DNA. RNA quality, assessed by RNA Integrity Number (RIN), were determined by Agilent Bioanalyzer (Agilent Technologies, Palo Alto, CA, USA). RNA samples of sufficient quality (RIN > 8) were used to construct RNA-seq libraries.

### RNAseq library preparation, sequencing and mapping

Library preparation and sequencing were done in the Penn State Genomics Core Facility (University Park, PA, USA). 200ng of RNA per sample were used as the input of TruSeq Stranded mRNA Library Prep kit to make barcoded libraries according to the manufacturer’s protocol (Illumina Inc., San Diego, CA, USA). The library concentration was determined by qPCR using the Library Quantification Kit Illumina Platforms (Kapa Biosystems, Boston, MA, USA). An equimolar pool of the barcoded libraries was made and the pool was sequencing on a HiSeq 2500 system (Illumina) in Rapid Run mode using single ended 150 base-pair reads. 30-37 million reads were acquired for each library. Mapping were done in the Penn State Bioinformatics Consulting Center. Quality trimming and adapter removal was performed using trimmomatic ^54^. The trimmed data were then mapped to the mouse genome (mm10, NCBI) using hisat2 alignment program. Hisat2 alignment was performed by specifying the ‘rna-strandness’ parameter. Default settings were used for all the other parameters ^55^. Coverages were obtained using bedtools ^56^. Mapped data was visualized and checked with IGV ^57^. Reads mapped to each gene were counted using featureCounts ^58^. The two RNAseq datasets reported in this article are available in NCBI Gene Expression Omnibus (GEO) under the accession number GSE143290.

### Data pre-processing

The raw count of *n*=24,421 transcripts was obtained after mapping in each of the two studies. Transcripts with low expression levels were first removed, for the purpose of assuring that each analysis would be focused on the tissue-specific mRNAs and would not be disrupted by the genes not expressed in that organ. In the SI study, if the expression level of a gene was lower than 10 in more than eight of the samples, or if the sum of the expression level in all 12 samples was lower than 200, this transcript was removed from the later analysis. In the colon study, if the expression level was lower than 10 in more than 10 of the samples, or if the sum of the expression level in all 16 samples was lower than 220, the transcript was regarded as lowly expressed gene and removed. As a result, 14,368 and 15,340 genes were remained in the SI and colon datasets, respectively. Both datasets, following normalization according to DESeq2 package, were used for the following DE analysis and co-expression network analysis (Fig. 2).

### Differential expression

Differential expression (DE) analysis were performed using the DESeq2 package ^59^. Because of the two-by-two factorial design, the effect of VA, *C. rodentium* infection and VA x *C. rodentium* Interaction were examined separately. This article focused on the VA effect, with adjusted *P*<0.05 (to ensure statistical significance) and |Fold change|> 2 (to ensure biological significance) as the criteria for differentially expressed genes (DEGs). To visualize the result and prioritize genes of interest, heatmaps and volcano plots were depicted using the R packages pheatmap, ggplot, and ggrepel.

### WGCNA

For consistency and comparability, the same data matrix in differential expression were used to build signed co-expression networks using WGCNA package in R ^60^. In other words, the dataset went through the same screening and normalization process as the differential expression analysis (Fig. 2). For each set of genes, a pair-wise correlation matrix was computed, and an adjacency matrix was calculated by raising the correlation matrix to a power. A parameter named Softpower was chosen using the scale-free topology criterion ^61^. An advantage of weighted correlation networks is the fact that the results are highly robust with respect to the choice of the power parameter. For each pairs of genes, a robust measure of network interconnectedness (topological overlap measure) was calculated based on the adjacency matrix. The topological overlap (TO) based dissimilarity was then used as input for average linkage hierarchical clustering. Finally, modules were defined as branches of the resulting clustering tree. To cut the branches, a hybrid dynamic tree-cutting algorithm was used ^62^. To obtain moderately large and distinct modules, we set the minimum module size to 30 genes and the minimum height for merging modules at 0.25. As per the standard of WGCNA tool, modules were referred to by their color labels henceforth and marked as SI(Color) or Colon(Color). Genes that did not belong to any modules were assigned to the Grey module. Color assignment together with hierarchical clustering dendrogram were depicted. To visualize the expression pattern of each module, heatmap were drawn using the pheatmap package in R. Eigengenes were computed and used as a representation of individual modules. Eigengenes are the first principal component of each set of module transcripts, describing most of the variance in the module gene expression. Eigengenes provides a coarse-grained representation of the entire module as a single entity ^20^. To identify modules of interest, module-trait relationships were analyzed. The modules were tested for their associations with the traits by correlating module eigengenes with trait measurements, i.e., categorical traits (VA status, infection status, and gender) and numerical traits (shedding, body weight, body weight change percentage, and RIN score). The correlation coefficient and significance of each correlation between module eigengenes and traits were visualized in a labeled heatmap, color-coded by blue (negative correlation) and red (positive correlation) shades. Because the present paper has a focus on the VA effect, the only trait reported was the VA status. The correlation results from all other traits were beyond the scope of this article and will be discussed elsewhere.

### Functional enrichment analyses

Gene ontology (GO) and KEGG enrichment were assessed using clusterProfiler (v3.6.0) in R ^63^. For DEG list and co-expression modules, the background was set as the total list of genes expressed in the SI (14,368 genes) or colon (15,340 genes), respectively. The statistical significance threshold level for all GO enrichment analyses was p.adj (Benjamini and Hochberg adjusted for multiple comparisons) <0.05.

### Cell type marker enrichment

The significance of cell type enrichment was assessed for each module in a given network with a cell type marker list of the same organ, using a one-sided hypergeometric test (as we were interested in testing for over-representation). To account for multiple comparisons, a Bonferroni correction was applied on the basis of the number of cell types in the organ-specific cell type marker list. The criterion for a module to significantly overlap with a cell marker list was set as a corrected hypergeometric *P* value < 0.05 ^22, 49, 64^. The cell type marker lists of SI and colon were obtained from CellMarker database (http://biocc.hrbmu.edu.cn/CellMarker/), a manually curated resource of cell markers ^21^.

### Module visualization

A package named igraph was used to visualize the co-expression network of genes in each individual modules based on the adjacency matrix computed by WGCNA. The connectivity of a gene is defined as the sum of connection strength (adjacency) with the other module members. In this way, the connectivity measures how correlated a gene is with all other genes in the same module ^65^. The top 50 genes in each module ranked by connectivity were used for the visualization. In order to further improve network visibility, adjacencies equal to 1 (adjacencies of a gene to itself) or lower than 0.7 (less important co-expression relationships) were not shown in the diagram. Each node represents a gene. The width of the lines between any two nodes was scaled based on the adjacency of this gene pair. The size of the node was set proportional to the connectivity of this gene. For the shape of the nodes, squares were used to represent all the mouse transcription factors (TFs), based on AnimalTFDB 3.0 database (http://bioinfo.life.hust.edu.cn/AnimalTFDB/<u>#!/</u>) ^66^. Circles represent all the non-TFs. In terms of the color of the nodes, upregulated DEGs in the VAS condition were marked with red color and downregulated DEGs in blue. For the font of the gene labels, red bold italic was used to highlight genes which are cell type markers according to CellMarker database (http://biocc.hrbmu.edu.cn/CellMarker/) ^21^.

### Module preservation

Genes with low expression (based on the criteria stated in Differential Expression session) in the SI or Colon datasets were excluded from this analysis. In other words, only the probes that were robustly expressed in both organs were used for module preservation analysis. Therefore, reconstruction of co-expression networks were done in both datasets separately, with minimum module size = 30 and the minimum height for merging modules = 0.25 ^60–62^. All the modules from this analysis were marked as mpSI(Color) or mpColon(Color).

Color assignment was remapped so that the corresponding modules in the SI network and the re-constructed mpSI network sharing the similar expression pattern were assigned with the same color. Similarly, the modules in the re-constructed mpColon network were also assigned same colors according to the expression pattern resemblance with their corresponding Colon modules. In order to quantify network preservation between mpSI and mpColon networks at the module level, permutation statistics is calculated in R function modulePreservation, which is included in the updated WGCNA package ^65^. Briefly, module labels in the test network (mpColon network) were randomly permuted for 50 times and the corresponding preservation statistics were computed, based on which Zsummary, a composite preservation statistic, were calculated. Zsummary combines multiple preservation Z statistics into a single overall measure of preservation ^65^. The higher the value of Zsummary, the more preserved the module is between the two datasets. As for threshold, if Zsummary >10, there is strong evidence that the module is preserved. If Zsummary is >2 but <10, there is weak to moderate evidence of preservation. If Zsummary <2, there is no evidence that the module is preserved ^65^. The Grey module contains uncharacterized genes while the Gold module contains random genes. In general, those two modules should have lower Z-scores than most of the other modules.

### Module overlap

Module overlap is the number of common genes between one module and a different module. The significance of module overlap can be measured using the hypergeometric test. In order to quantify module comparison, in other words, to test if any of the modules in the test network (mpColon network) significantly overlaps with the modules in the reference network (mpSI network), the overlap between all possible pairs of modules was calculated along with the probability of observing such overlaps by chance. The significance of module overlap was assessed for each module in a given network with all modules in the comparison network using a one-sided hypergeometric test (as we were interested in testing for over-representation) ^49^. To account for multiple comparisons, a Bonferroni correction was applied on the basis of the number of modules in the comparison network (mpColon network). Module pair with corrected hypergeometric *P* value < 0.05 was considered a significant overlap.

## Acknowledgements

We would like to acknowledge the sequencing service performed by C. Praul and colleagues in the Genomic Core Facility at the Pennsylvania State University. The study was supported by research funding (R56 AI114972 and R01 GM109453) and training grant (T32GM108563) provided by National Institute of Health (NIH) as well as J. Lloyd Huck Graduate Fellowship and Dorothy Foehr Huck Chair endowment provided by the Huck Institutes of Life Sciences. The funders had no role in the study design, data collection and analysis, decision to publish, or preparation the manuscript.

## Author Contributions

A.C.R., M.T.C., and Z.C. conceived the study, designed the experiments, and made major contributions to drafting the manuscript. A.C.R. and M.T.C. supervised the projects and organized the coworkers. Z.C. performed most of the experiments and data analyses. Y.L. and Q.L. provided guidance on the selection and application of bioinformatics tools. C.W. measured the serum retinol level using UPLC and Q.C., L.M.S., and V.W. assisted with the animal studies. A.S. and I.A. contributed on mapping, data pre-processing, and bioinformatics consulting. All authors reviewed the text and approved the final manuscript.

## Tables

The only tables are included as supplementary material.

## Data availability

The RNAseq datasets generated for this study can be found in NCBI Gene Expression Omnibus (GEO) database with the accession number GSE143290 at https://www.ncbi.nlm.nih.gov/geo/query/acc.cgi?acc=GSE143290.

---

**Supplementary Figure 1.**
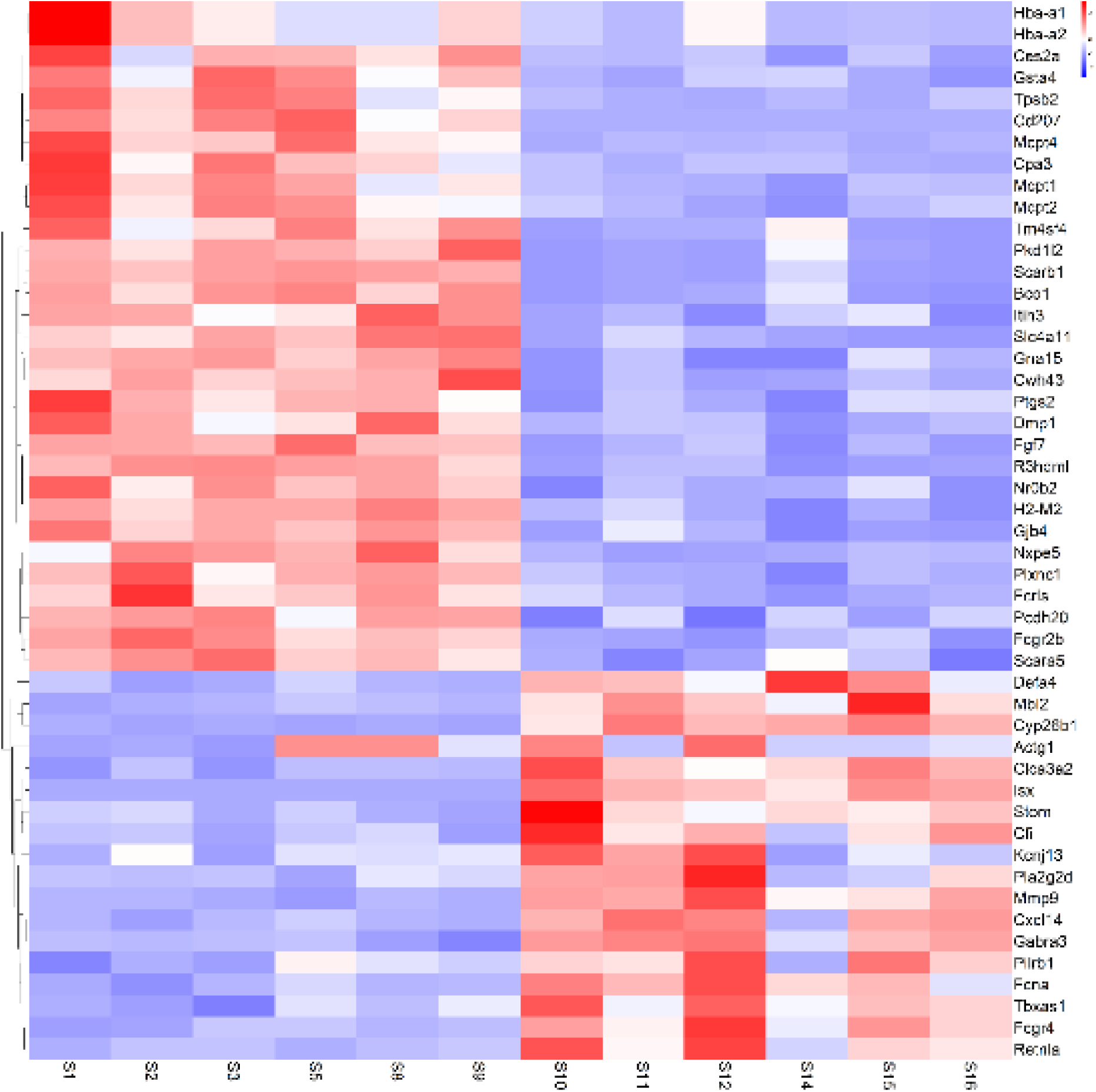
Heat map for DEGs corresponding to VA effect in the SI study. **Notes:** Red: group mean expression significantly higher; Blue: group mean expression significantly lower. VAD: columns S1, S2, S3, S5, S8, and S9. VAS: columns S10, S11, S12, S14, S15, and S16. **Abbreviations:** differentially expressed gene (DEG), vitamin A deficient (VAD), vitamin A sufficient (VAS).

**Supplementary Figure 2.**
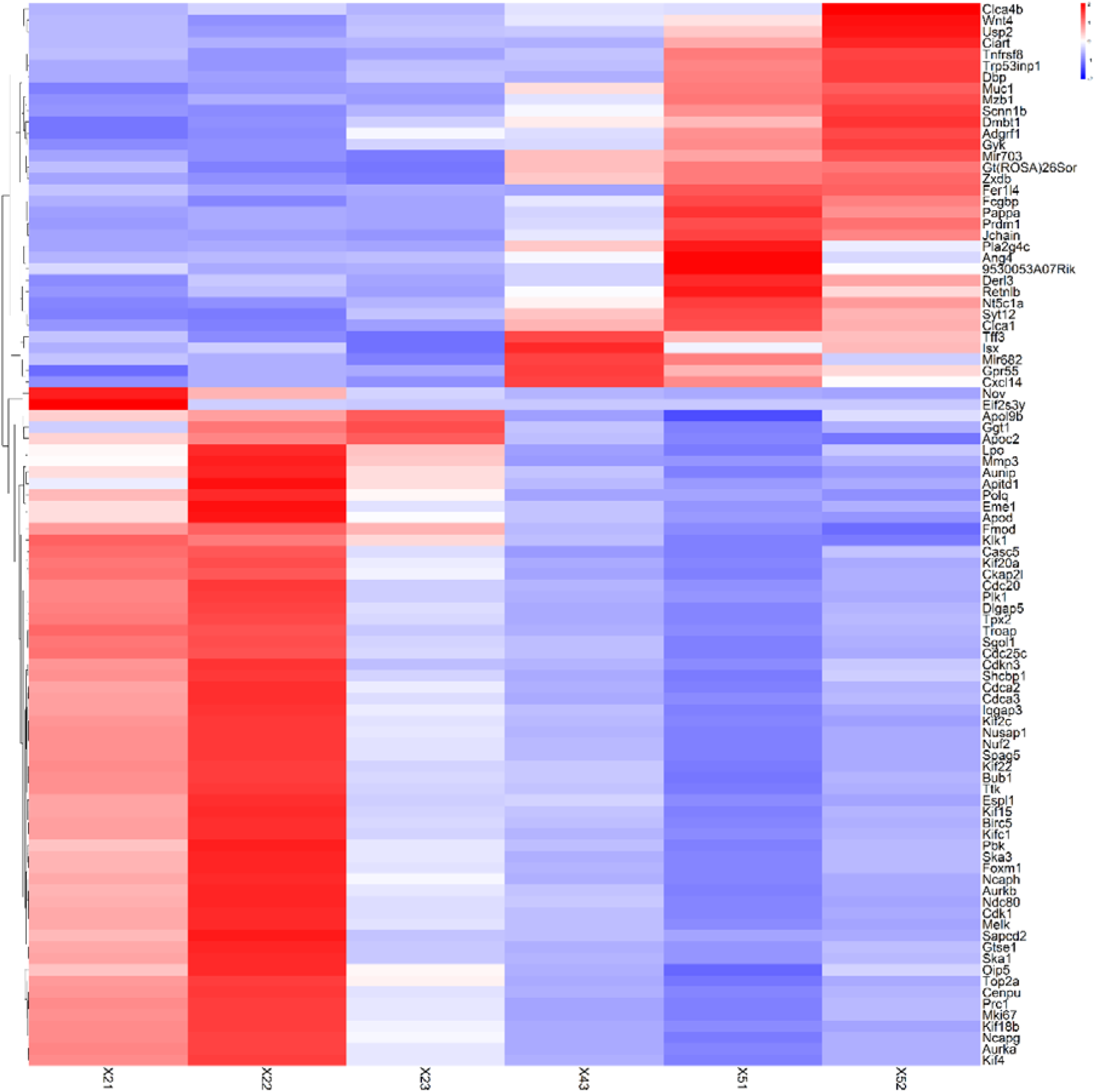
Heat map for DEGs corresponding to VA effect in the colon study. **Notes:** Red: group mean expression significantly higher; Blue: group mean expression significantly lower. VAD: columns X21, X22 and X23. VAS: columns X43, X51 and X52. **Abbreviations:** differentially expressed gene (DEG), vitamin A deficient (VAD), vitamin A sufficient (VAS).

**Supplementary Figure 3.**
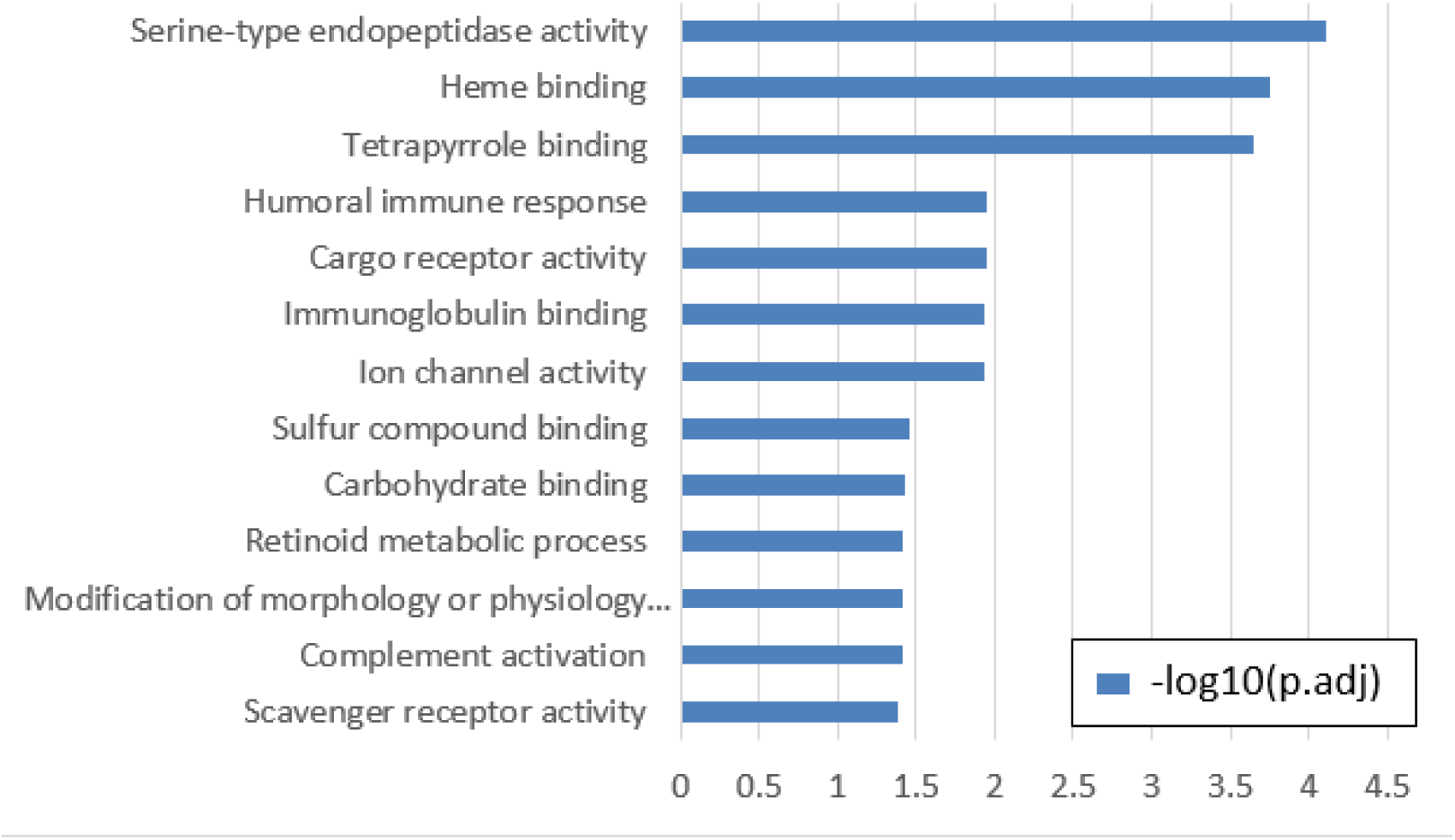
GO enrichment terms of DEGs corresponding to VA effect in the SI study. Criteria: p.adj<0.05. **Abbreviations:** p.adj (BH-adjusted p value from the hypergeometric test conducted by ClusterProfiler), Gene ontology (GO), differentially expressed gene (DEG)

**Supplementary Figure 4.**
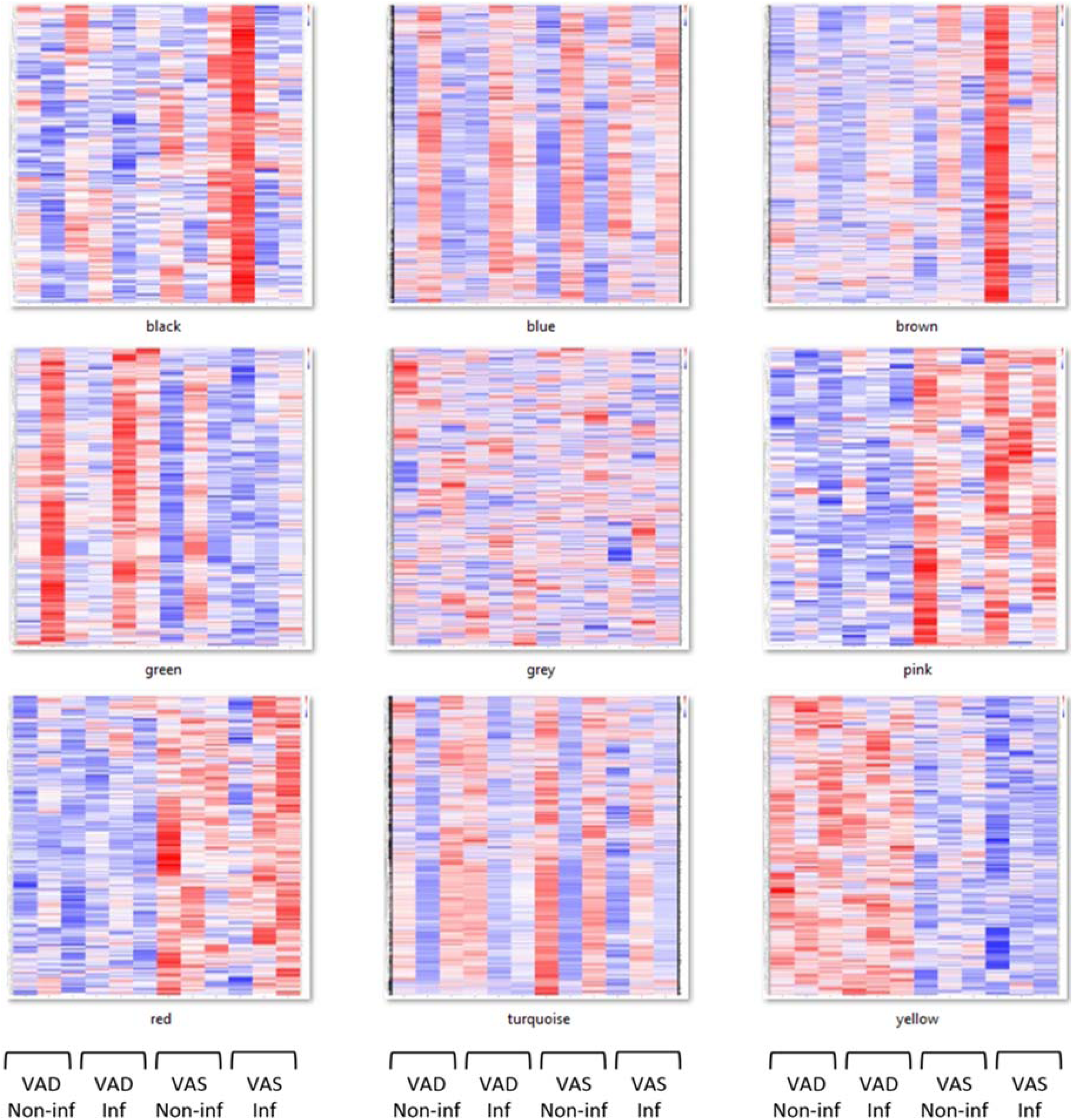
Gene co-expression patterns identified by WGCNA in the SI study. Notes: Red: High expression level; Blue: Low expression level. VAD: columns 1, 2, 3, 4, 5, and 6. VAS: columns 7, 8, 9, 10, 11, and 12. **Abbreviations:** vitamin A deficient (VAD), vitamin A sufficient (VAS), weighted gene co-expression network analysis (WGCNA).

**Supplementary Figure 5.**
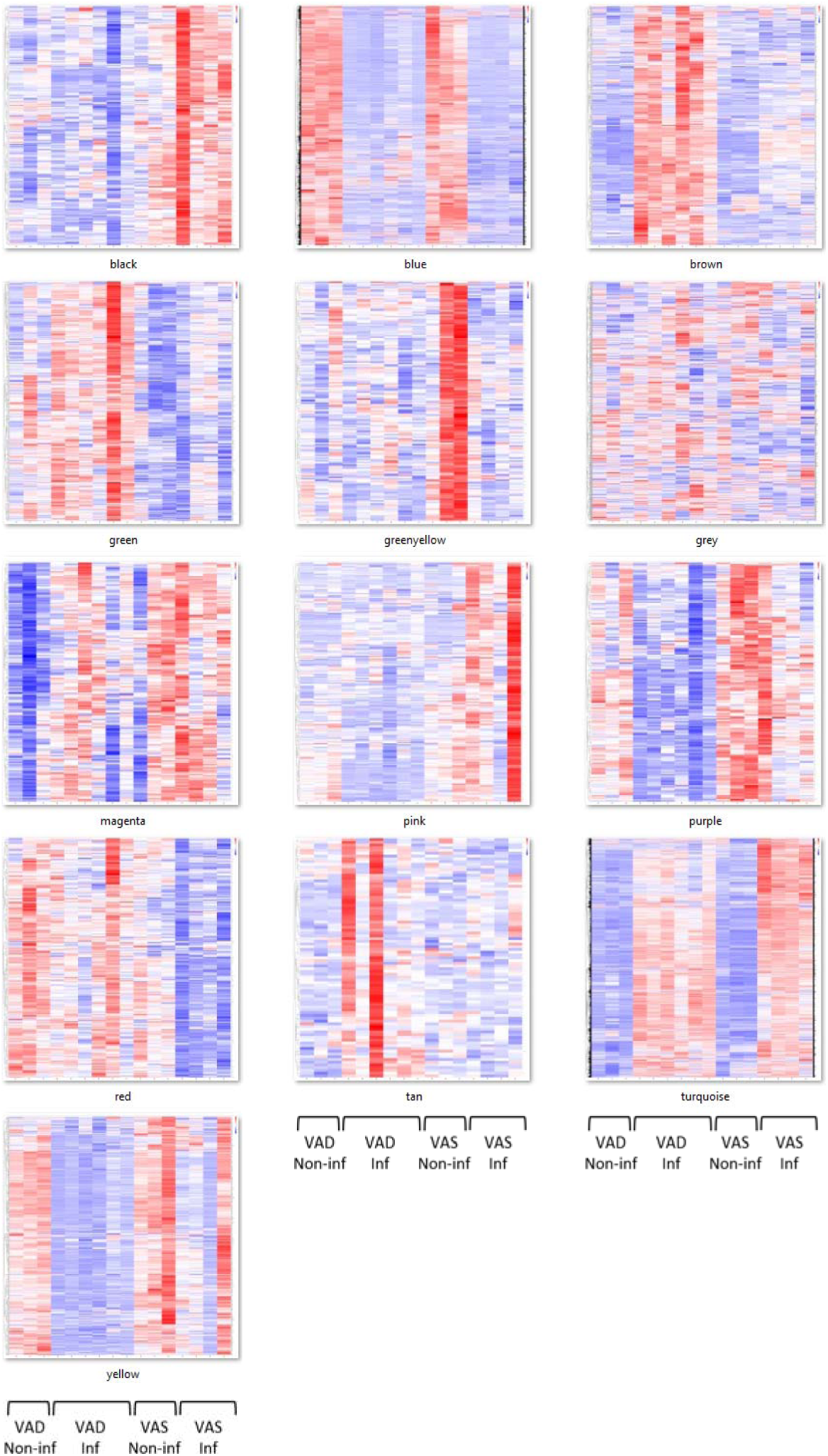
Gene co-expression patterns identified by WGCNA in the colon study. **Notes:** Red: High expression level; Blue: Low expression level. VAD: columns 1, 2, 3, 4, 5, 6, 7, 8, and 9. VAS: columns 10, 11, 12, 13, 14, 15, and 16. **Abbreviations:** vitamin A deficient (VAD), vitamin A sufficient (VAS), weighted gene co-expression network analysis (WGCNA).

**Supplementary Figure 6.**
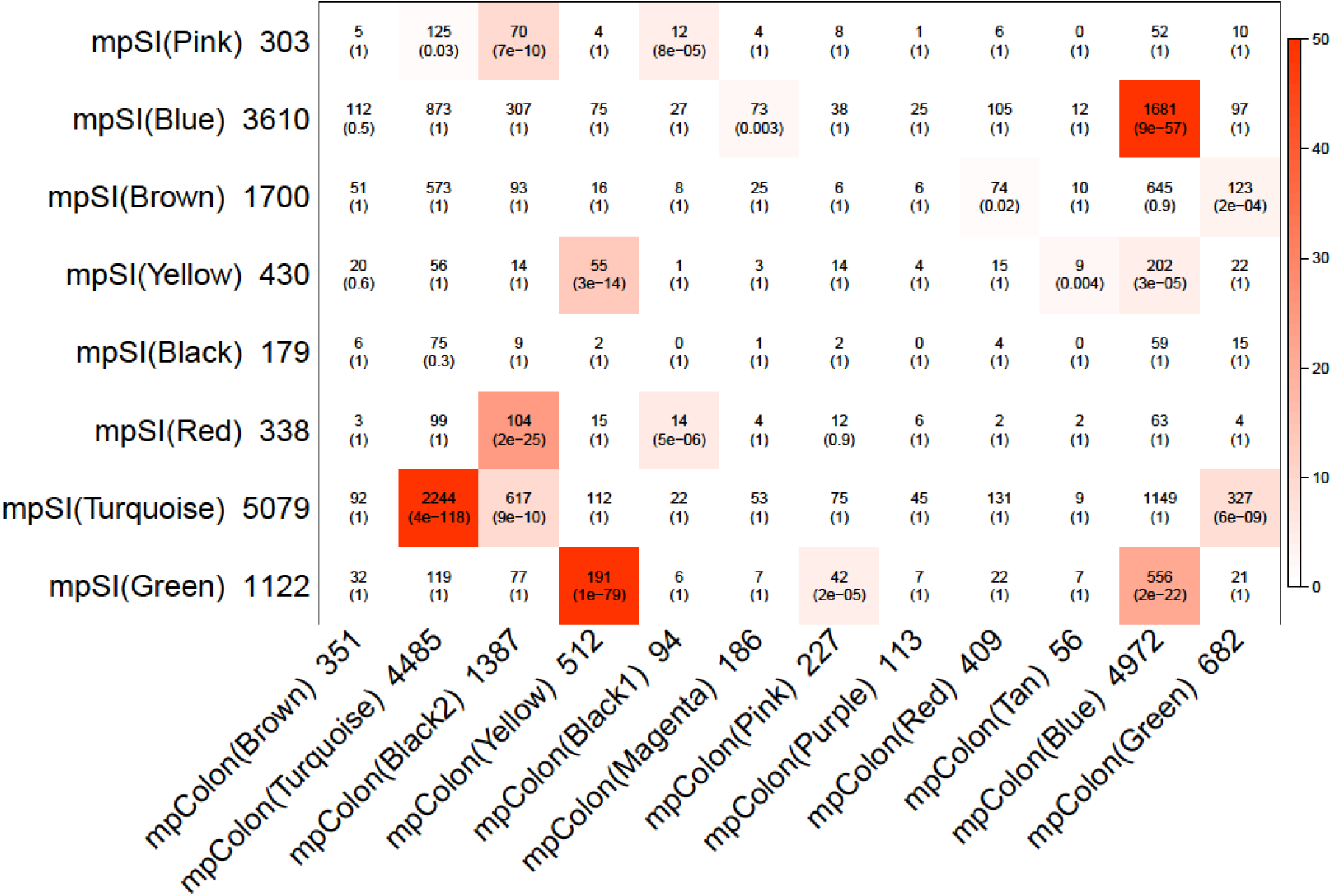
Cross-tabulation of SI modules (rows) and colon modules (columns). **Notes:** Each row and column is labeled by (color assignment module size). In each cell of the table, upper numbers give counts of genes in the intersection of the corresponding row and column modules. Lower numbers in parentheses: Bonferroni corrected p-value (CP) of the one-sided hypergeometric test, which evaluates if a significant overlap is reached between the corresponding modules. Criteria: CP<0.05. The table is color-coded by –log10(CP), according to the color legend on the right.

**Supplementary Table 1.** Upregulated DEGs in the VAS group compared with VAD group in the SI study. The order is ranked by the adjusted p values (padj). **Abbreviations:** BaseMean: Mean expression level (normalized by DESeq2) across all 12 libraries Fold Change: Normalized mean expression level of VAS group / Normalized mean expression level of VAD group padj: adjusted p value calculated by DESeq2

|  | BaseMean | Fold Change | padj | EntrezID | Fullname |
| --- | --- | --- | --- | --- | --- |
| Isx | 957.15 | 3231.03 | 4.92E-24 | 71597 | intestine specific homeobox |
| Cyp26b1 | 39.63 | 14.12 | 1.52E-12 | 232174 | cytochrome P450, family 26, subfamily b, polypeptide 1 |
| Mmp9 | 53.98 | 10.06 | 1.60E-09 | 17395 | matrix metalloproteinase 9 |
| Mbl2 | 164.18 | 10.37 | 2.02E-09 | 17195 | mannose-binding lectin (protein C) 2 |
| Retnla | 32.95 | 7.93 | 0.000147 | 57262 | resistin like alpha |
| Clca3a2 | 464.34 | 2.22 | 0.000235 | 80797 | chloride channel accessory 3A2 |
| Fcgr4 | 81.24 | 3.17 | 0.000337 | 246256 | Fc receptor, IgG, low affinity IV |
| Pla2g2d | 58.32 | 3.13 | 0.002081 | 18782 | phospholipase A2, group IID |
| Gabra3 | 32.47 | 3.18 | 0.002919 | 14396 | gamma-aminobutyric acid (GABA) A receptor, subunit alpha 3 |
| Kcnj13 | 105.76 | 2.18 | 0.003246 | 100040591 | potassium inwardly-rectifying channel, subfamily J, member 13 |
| Cxcl14 | 400.56 | 2.08 | 0.003349 | 57266 | chemokine (C-X-C motif) ligand 14 |
| Fcna | 30.21 | 2.67 | 0.004892 | 14133 | ficolin A |
| Tbxas1 | 50.77 | 2.46 | 0.00581 | 21391 | thromboxane A synthase 1, platelet |
| Pilrb1 | 36.20 | 3.63 | 0.013114 | 170741 | paired immunoglobulin-like type 2 receptor beta 1 |
| Cfi | 106.97 | 3.33 | 0.014409 | 12630 | complement component factor i |
| Actg1 | 42795.18 | 2.04 | 0.030234 | 11465 | actin, gamma, cytoplasmic 1 |
| Defa4 | 661.50 | 2.08 | 0.042915 | 13238 | defensin, alpha 4 |
| Stom | 6202.03 | 2.09 | 0.045444 | 13830 | stomatin |

**Supplementary Table 2.** Downregulated DEGs in the VAS group compared with VAD group in the SI study. The order is ranked by the adjusted p values (padj). **Abbreviations:** BaseMean: Mean expression level (normalized by DESeq2) across all 12 libraries Fold Change: -Normalized mean expression level of VAD group/ Normalized mean expression level of VAS group padj: adjusted p value calculated by DESeq2

|  | BaseMean | Fold Change | padj | EntrezID | Fullname |
| --- | --- | --- | --- | --- | --- |
| Cd207 | 90.13 | -1055.17 | 7.41E-12 | 246278 | CD207 antigen |
| Fcgr2b | 197.54 | -2.17 | 1.95E-08 | 14130 | Fc receptor, IgG, low affinity IIb |
| Mcpt4 | 42.09 | -18.32 | 3.06E-08 | 17227 | mast cell protease 4 |
| R3hdml | 41.80 | -3.91 | 3.01E-07 | 1E+08 | R3H domain containing-like |
| Fgf7 | 125.51 | -2.44 | 3.78E-05 | 14178 | fibroblast growth factor 7 |
| Tpsb2 | 20.66 | -8.61 | 0.000147 | 17229 | tryptase beta 2 |
| Cwh43 | 47.30 | -3.56 | 0.000159 | 231293 | cell wall biogenesis 43 C-terminal homolog |
| Scarb1 | 15409.55 | -18.32 | 0.000173 | 20778 | scavenger receptor class B, member 1 |
| H2-M2 | 195.61 | -2.14 | 0.000579 | 14990 | histocompatibility 2, M region, locus 2 |
| Cpa3 | 25.50 | -5.29 | 0.001197 | 12873 | carboxypeptidase A3, mast cell |
| Mcpt1 | 201.78 | -5.57 | 0.001313 | 17224 | mast cell protease 1 |
| Slc4a11 | 73.43 | -2.35 | 0.002602 | 269356 | solute carrier family 4 sodium bicarbonate, transporter-like, member 11 |
| Nr0b2 | 46.58 | -3.60 | 0.002602 | 23957 | nuclear receptor subfamily 0, group B, member 2 |
| Scara5 | 64.88 | -2.35 | 0.003349 | 71145 | scavenger receptor class A member 5 |
| Fcrls | 75.39 | -2.69 | 0.004543 | 80891 | Fc receptor-like S, scavenger receptor |
| Gsta4 | 2001.56 | -3.71 | 0.004722 | 14860 | glutathione S-transferase, alpha 4 |
| Pkd1l2 | 82.38 | -17.15 | 0.00499 | 76645 | polycystic kidney disease 1 like 2 |
| Plxnc1 | 92.51 | -2.19 | 0.00581 | 54712 | plexin C1 |
| Nxpe5 | 38.78 | -2.74 | 0.00581 | 381680 | neurexophilin and PC-esterase domain family, member 5 |
| Mcpt2 | 79.94 | -4.45 | 0.006874 | 17225 | mast cell protease 2 |
| Ptgs2 | 184.93 | -2.35 | 0.00872 | 19225 | prostaglandin-endoperoxide synthase 2 |
| Pcdh20 | 75.28 | -2.13 | 0.010642 | 219257 | protocadherin 20 |
| Tm4sf4 | 5881.59 | -4.07 | 0.010642 | 229302 | transmembrane 4 superfamily member 4 |
| Bco1 | 18783.84 | -8.66 | 0.017006 | 63857 | beta-carotene oxygenase 1 |
| Itih3 | 56.45 | -2.50 | 0.029795 | 16426 | inter-alpha trypsin inhibitor, heavy chain 3 |
| Dmp1 | 174.24 | -3.49 | 0.030234 | 13406 | dentin matrix protein 1 |
| Gna15 | 32.73 | -2.33 | 0.032603 | 14676 | guanine nucleotide binding protein,<br>alpha 15 |
| Hba-a1 | 1116.80 | -3.15 | 0.042345 | 15122 | hemoglobin alpha, adult chain 1 |
| Hba-a2 | 1116.81 | -3.15 | 0.042345 | 110257 | hemoglobin alpha, adult chain 2 |
| Ces2a | 7062.11 | -2.19 | 0.044776 | 102022 | carboxylesterase 2A |
| Gjb4 | 18.73 | -3.44 | 0.049916 | 14621 | gap junction protein, beta 4 |

**Supplementary Table 3.** Upregulated DEGs in the VAS group compared with VAD group in the colon study. The order is ranked by the adjusted p values (padj). **Abbreviations:** BaseMean: Mean expression level (normalized by DESeq2) across all 16 libraries Fold Change: Normalized mean expression level of VAS group/ Normalized mean expression level of VAD group padj: adjusted p value calculated by DESeq2

|  | BaseMean | Fold change | padj | EntrezID | Fullname |
| --- | --- | --- | --- | --- | --- |
| Pla2g4c | 153.87 | 11.61 | 1.77E-06 | 232889 | phospholipase A2 group IVC (cytosolic calcium-independent) |
| Tff3 | 14507.68 | 2.31 | 0.000218 | 21786 | trefoil factor 3 intestinal |
| Muc1 | 1208.43 | 2.84 | 0.000576 | 17829 | mucin 1 |
| Retnlb | 4272.92 | 8.44 | 0.000801 | 57263 | resistin like beta |
| Gyk | 5346.19 | 2.19 | 0.001731 | NA | NA |
| Jchain | 8052.94 | 5.80 | 0.001902 | 16069 | immunoglobulin joining chain |
| Ciart | 70.23 | 7.68 | 0.002115 | 229599 | circadian associated repressor of transcription |
| Fcgbp | 58520.79 | 2.14 | 0.002649 | 215384 | Fc fragment of IgG binding protein |
| Isx | 478.87 | 3.75 | 0.003429 | 71597 | intestine specific homeobox |
| Ang4 | 737.71 | 7.73 | 0.003836 | 219033 | Angiogenin ribonuclease A family member 4 |
| Zxdb | 128.03 | 2.29 | 0.004079 | 668166 | zinc finger X-linked duplicated B |
| Tnfrsf8 | 253.70 | 3.13 | 0.004473 | 21941 | tumor necrosis factor receptor superfamily member 8 |
| Dmbt1 | 256019.57 | 2.05 | 0.006472 | 12945 | deleted in malignant brain tumors 1 |
| Prdm1 | 365.97 | 2.46 | 0.007372 | 12142 | PR domain containing 1 with ZNF domain |
| Gpr55 | 62.18 | 2.39 | 0.007529 | 227326 | G protein-coupled receptor 55 |
| Scnn1b | 171.80 | 3.36 | 0.007837 | 20277 | sodium channel nonvoltage-gated 1 beta |
| Cxcl14 | 237.56 | 2.27 | 0.008616 | 57266 | chemokine (C-X-C motif) ligand 14 |
| Gt(ROSA)26Sor | 640.78 | 2.57 | 0.009265 | 14910 | gene trap ROSA 26 Philippe Soriano |
| Fer1l4 | 1111.67 | 3.67 | 0.010737 | 74562 | fer-1-like 4 (C. elegans) |
| Mzb1 | 499.91 | 2.70 | 0.014338 | 69816 | marginal zone B and B1 cell-specific protein 1 |
| Wnt4 | 153.06 | 2.76 | 0.014572 | 22417 | wingless-type MMTV integration site family member 4 |
| Dbp | 598.43 | 6.36 | 0.015664 | 13170 | D site albumin promoter binding protein |
| Adgrf1 | 1333.45 | 2.60 | 0.015726 | 77596 | adhesion G protein-coupled receptor F1 |
| Nt5c1a | 241.89 | 2.69 | 0.018873 | 230718 | 5'-nucleotidase cytosolic IA |
| Derl3 | 132.87 | 2.79 | 0.019995 | 70377 | Der1-like domain family member 3 |
| Clca4b | 35258.66 | 4.52 | 0.022243 | 99709 | chloride channel accessory 4B |
| Syt12 | 117.12 | 2.39 | 0.030138 | 171180 | synaptotagmin XII |
| Mir703 | 311.12 | 2.70 | 0.030376 | 735265 | microRNA 703 |
| 9530053A07Rik | 150.87 | 3.94 | 0.037762 | 319482 | RIKEN cDNA 9530053A07 gene |
| Clca1 | 33895.13 | 2.76 | 0.041848 | 23844 | chloride channel accessory 1 |
| Trp53inp1 | 2245.74 | 2.04 | 0.042511 | 60599 | transformation related protein 53 inducible nuclear protein 1 |
| Pappa | 95.62 | 3.85 | 0.043324 | 18491 | pregnancy-associated plasma protein A |
| Mir682 | 1134.07 | 2.44 | 0.044732 | 751556 | microRNA 682 |
| Usp2 | 574.80 | 2.10 | 0.045265 | 53376 | ubiquitin specific peptidase 2 |

**Supplementary Table 4.**
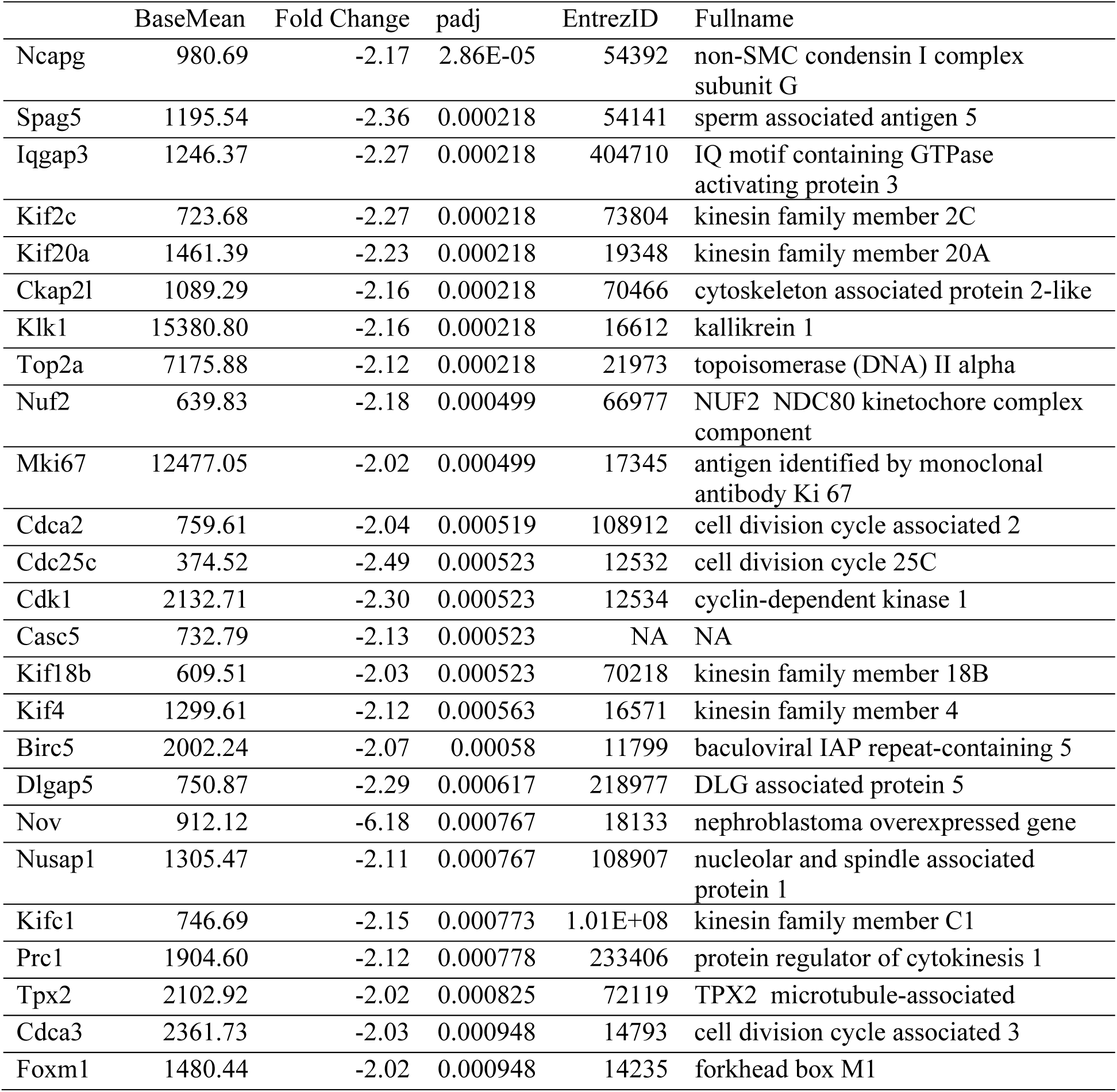
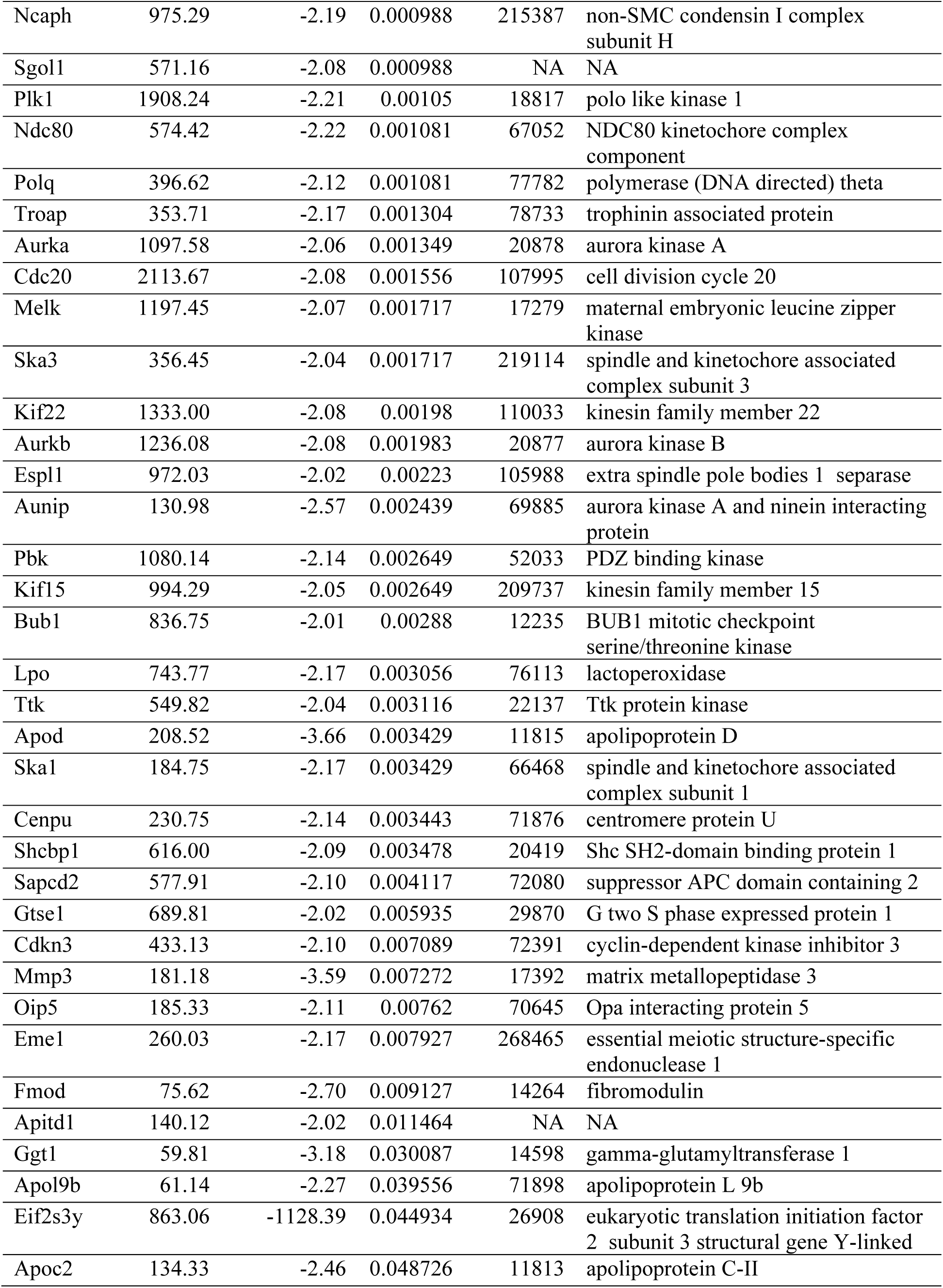
Downregulated DEGs in the VAS group compared with VAD group in the colon study. The order is ranked by the adjusted p values (padj). **Abbreviations:** baseMean: Mean expression level (normalized by DESeq2) across all 16 libraries Fold Change: -Normalized mean expression level of VAD group / Normalized mean expression level of VAS group padj: adjusted p value calculated by DESeq2

**Supplementary Table 5.**
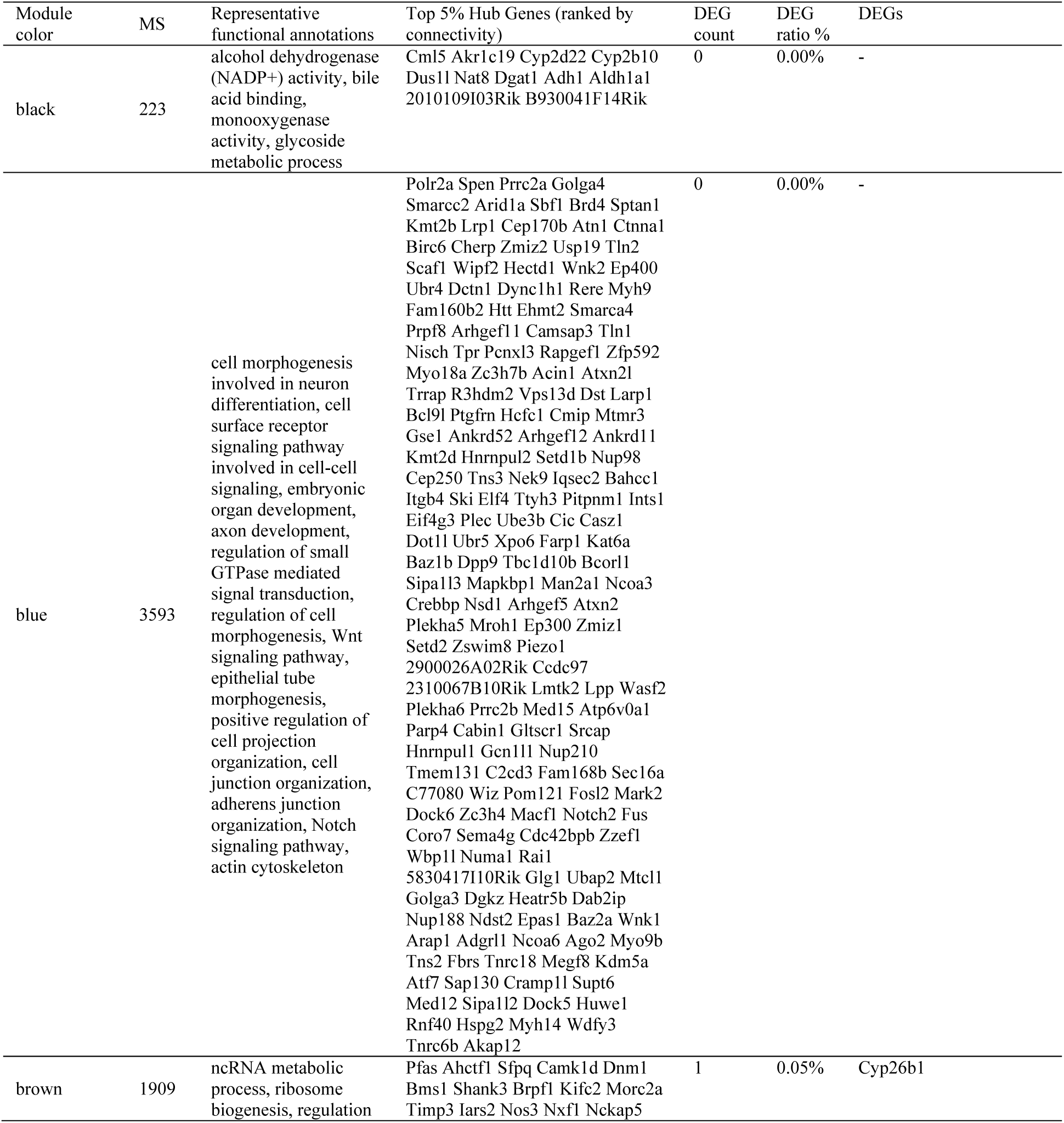
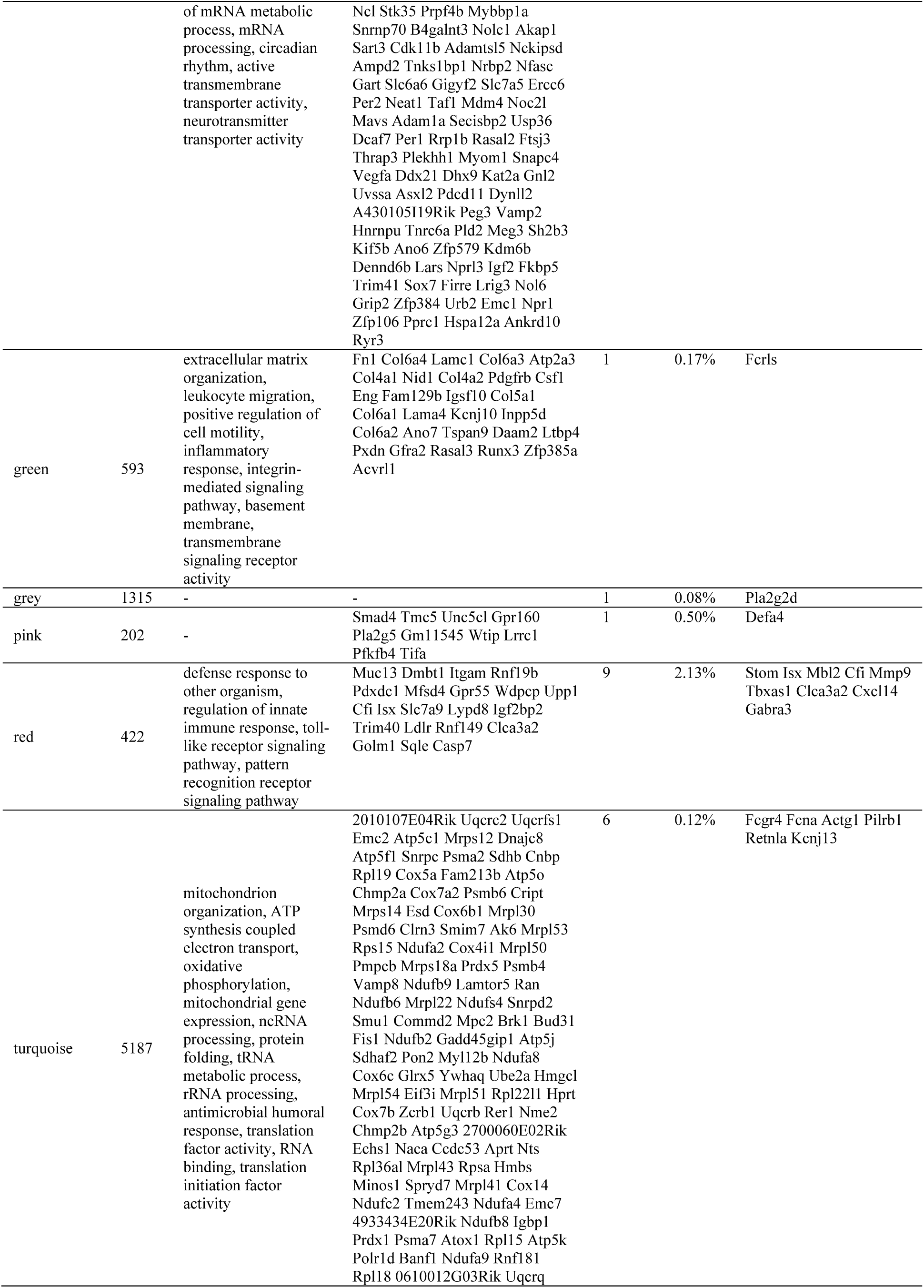
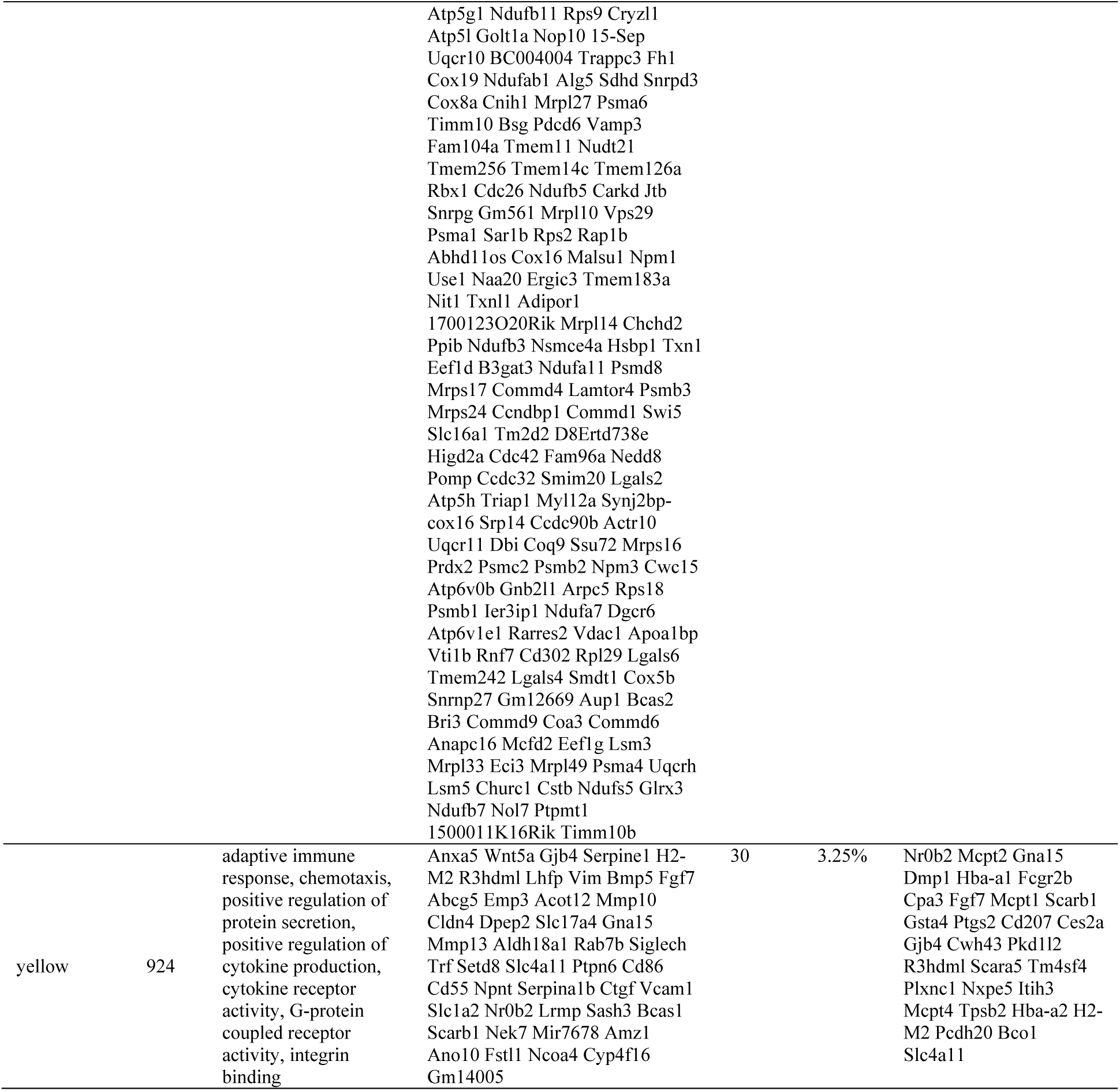
SI Modules and their sizes (MS), functional annotations, hub genes, and overlap with DEGs. **Abbreviations:** MS: Module Size. Number of gene in certain modules. DEG count: number of DEGs (corresponding to VA effect) that also belong to certain module DEG ratio: DEG count/MS

**Supplementary Table 6.**
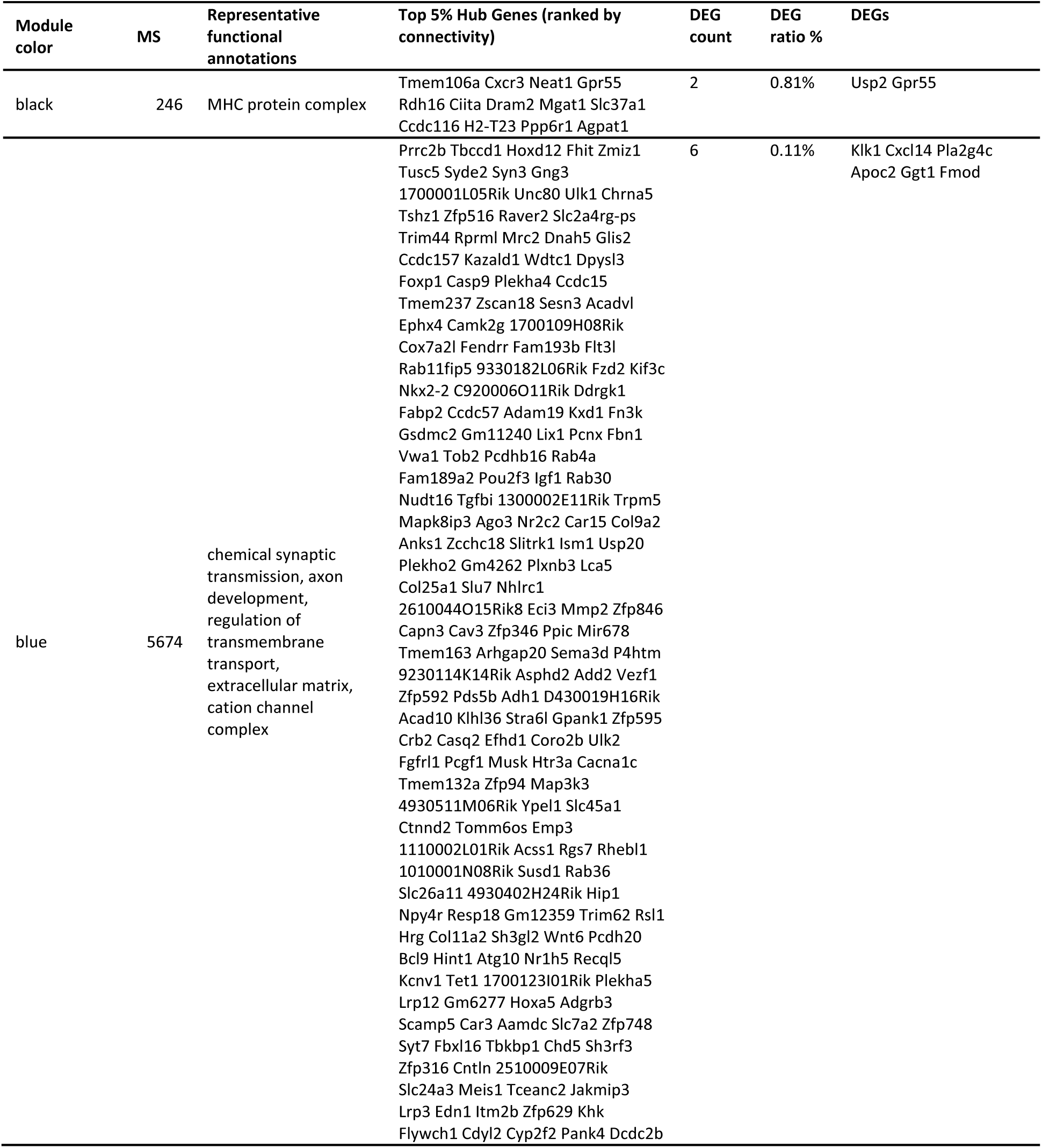
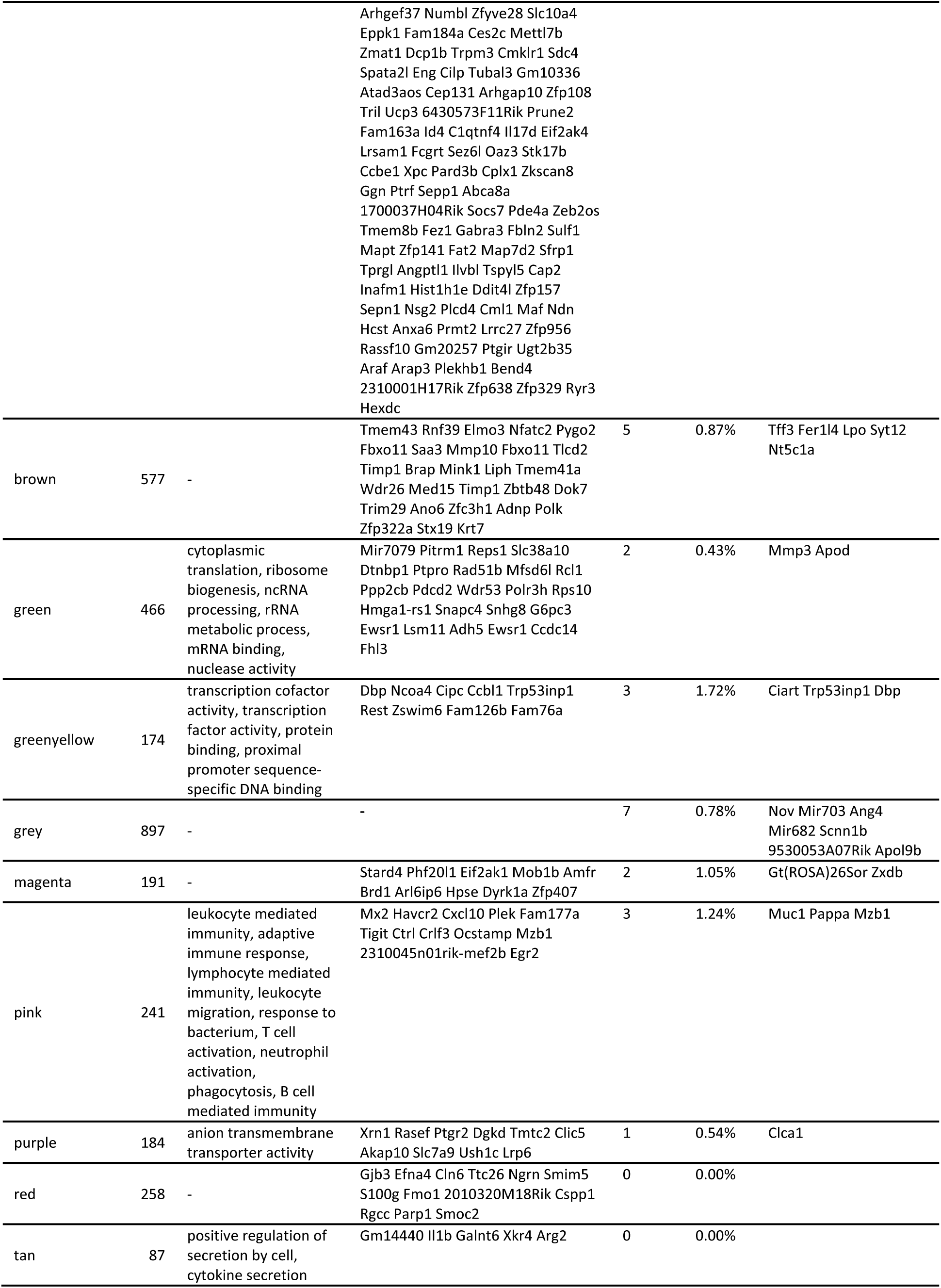
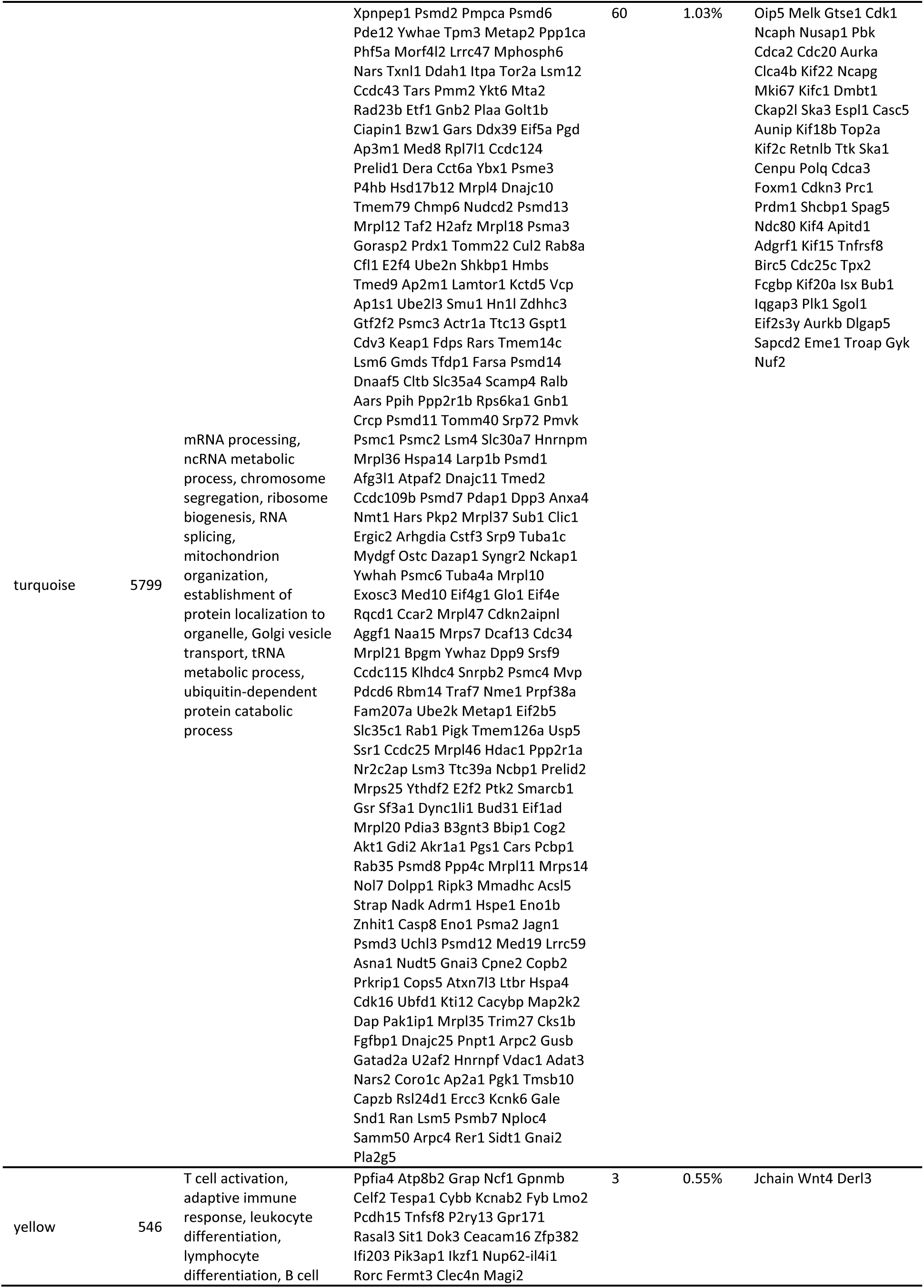
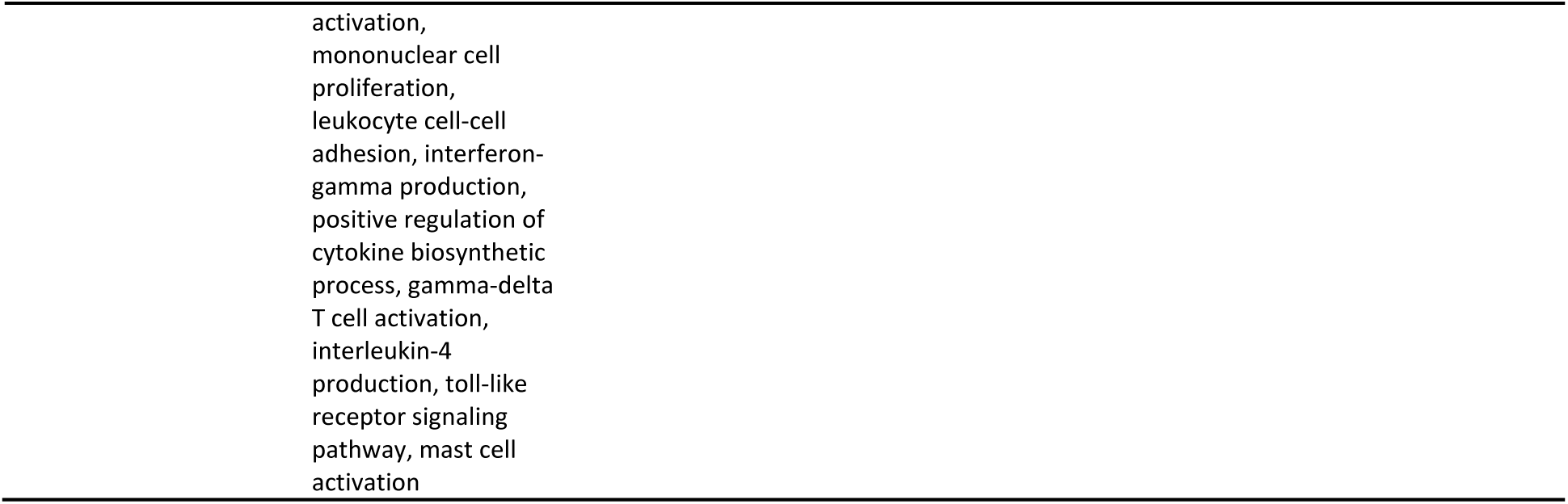
Colon Modules and their sizes (MS), functional annotations, hub genes, and overlap with DEGs. **Abbreviations:** MS: Module Size. Number of gene in certain modules. DEG count: number of DEGs (corresponding to VA effect) that also belong to certain module DEG ratio: DEG count/MS

**Supplementary Table 7.** Preservation of SI modules according to Zsummary. **Notes:** if Zsummary>10, there is strong evidence that the module is preserved. If Zsummary>2 but Zsummary<10, there is weak to moderate evidence of preservation. If Zsummary<2, there is no evidence that the module preserved. The Grey module contains uncharacterized genes while the Gold module contains random genes. The table is ranked by Zsummary, from high to low.

| Module color | Zsummary | Evidence of preservation |
| --- | --- | --- |
| Green | 13.546 | Strong |
| Turquoise | 11.471 | Strong |
| Red | 8.554 | Weak to moderate |
| Gold | 7.746 | Weak to moderate |
| Pink | 7.275 | Weak to moderate |
| Blue | 6.287 | Weak to moderate |
| Yellow | 3.519 | Weak to moderate |
| Brown | 2.544 | Weak to moderate |
| Black | 2.001 | Weak to moderate |
| Grey | -0.619 | No |

